# Acetylation of NDUFA13 at K7 Senses OXPHOS Dysfunction to Disrupt Prohibitin Scaffolds and Drive Cristae Remodelling in Heart Failure

**DOI:** 10.64898/2026.10.26.761921

**Authors:** Xinnuo Yang, Yinuo Lin, Jian Shen, Hongpeng Chen, Zihan Xu, Jian Ye, Yaping Wang, Danyang Xiong, Changle Ke, Lianlian Zhu, Xiangmin Kong, Zhiwei Zhong, Tanchen Ren, Jun Jiang, Xuwen Wang, Binben Wang, Duoduo Zhao, Zheng-zheng Li, Zhehua Shao, Bingchen Liu, Xiao He, Jinliang Nan

## Abstract

Mitochondrial cristae disintegration is a hallmark of heart failure, yet how oxidative phosphorylation (OXPHOS) dysfunction triggers this structural collapse remains unclear. Here we identify acetylation of the complex I subunit NDUFA13 at lysine 7 (K7) as a metabolic sensor coupling OXPHOS dysfunction to cristae remodelling. OXPHOS inhibition drove NDUFA13-K7 hyperacetylation, inducing conformational compaction and translocation to the intermembrane space. Acetylated NDUFA13 competitively engaged PHB2 through its coiled-coil domain, displacing PHB1 and disassembling prohibitin scaffolds. The acetylation-mimetic NDUFA13-K7Q induced cristae fragmentation *in vitro*, and cardiomyocyte-specific K7Q knock-in mice showed accelerated mortality and cristae loss under pressure overload, accompanied by impaired TCA cycle flux, defective fatty acid oxidation and selective depletion of polyunsaturated cardiolipins. These findings establish NDUFA13-K7 acetylation as a reversible switch converting bioenergetic failure into pathological cristae remodelling and metabolic reprogramming, highlighting therapeutic potential for targeted deacetylation.

**Graphical Abstract:** 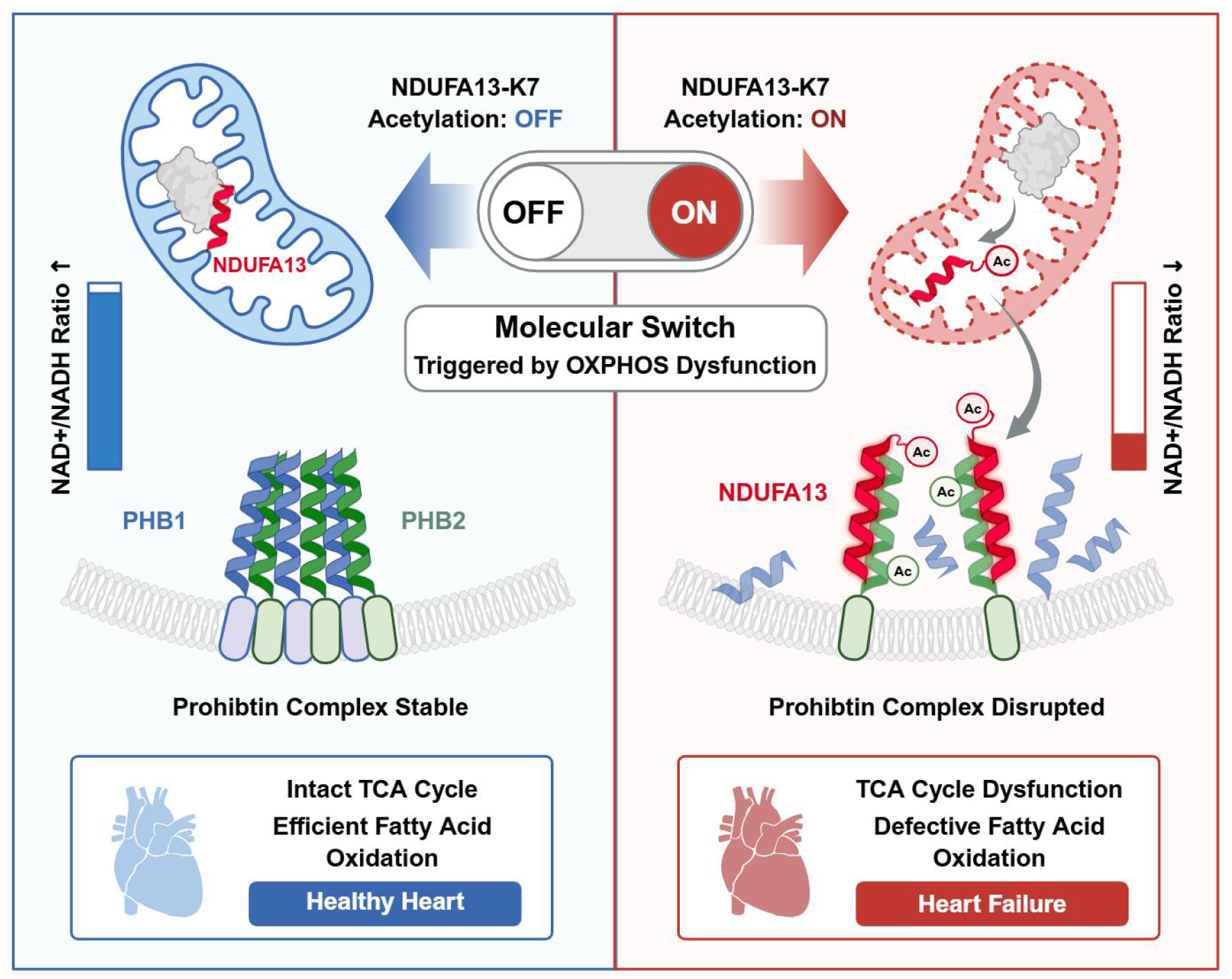

## Introduction

Heart failure (HF) is a leading cause of mortality worldwide, affecting more than 64 million people globally^1–3^. Impaired cardiac energy metabolism represents a core pathophysiological feature of this syndrome. Mitochondria occupy approximately 30% of cardiomyocyte volume and serve as the primary site for ATP production. In the healthy heart, mitochondria exhibit highly ordered cristae structures that maximise the efficiency of oxidative phosphorylation (OXPHOS)^4,5^. However, clinical studies consistently demonstrate that terminally failing hearts exhibit a markedly reduced density of cristae and disordered morphology, structural disruptions that are closely associated with diminished OXPHOS capacity, impaired substrate oxidation, deteriorating cardiac function, and a poor prognosis^6–8^. Although cristae remodelling is well established as a hallmark of HF, the molecular mechanisms by which OXPHOS dysfunction actively initiates the disintegration of cristae have yet to be fully elucidated.

The prohibitin (PHB) complex, composed of PHB1 and PHB2, serves as a critical scaffold protein for maintaining morphology of the inner mitochondrial membrane of the cristae^9,10^. Recent cryo-ET studies have revealed that PHB1/2 is assembled as a dodecamer of PHB1-PHB2 heterodimers anchored to the inner membrane, forming parallel interactions through coiled-coil domains that stabilize cristae curvature^11^. In both animal models and patient samples of HF, PHB1/2 and its higher-order super-complex (PHB1/2-SC) assembly are markedly reduced, accompanied by aberrant OPA1 cleavage and reduced cristae density^12,13^. However, the mechanism by which OXPHOS dysfunction triggers PHB1/2 disassembly remains unknown.

A key metabolic consequence of OXPHOS inhibition is a decline in the NAD^+^/NADH ratio, leading to reduced activity of deacetylases such as SIRT3 and consequent widespread hyperacetylation of mitochondrial proteins^14^. The acetylation levels of mitochondrial protein are significantly elevated in failing hearts^15^; however, the critical sites that mediate signal transduction from metabolic disruption to structural collapse have yet to be identified. As the rate-limiting step of OXPHOS and the primary site of NADH oxidation, whether accessory subunits of respiratory chain complex I can sense NAD^+^ changes and relay signals to cristae scaffolds has yet to be investigated^16,17^.

In this study, we aimed to elucidate how OXPHOS dysfunction actively drives the pathological remodelling of cristae. We hypothesised that the respiratory chain itself may include as yet unidentified ‘sentinel’ molecules that are capable of directly sensing metabolic stress and transmitting signals to scaffold proteins in the cristae. By integrating human HF sample analysis, unbiased proteomic screening, molecular dynamics simulations, and cardiomyocyte-specific knock-in mouse models, we discovered that the complex I accessory subunit NDUFA13 directly couples OXPHOS dysfunction to PHB1/2 scaffold disassembly *via* K7 acetylation-mediated protein–protein interactions. This finding establishes a novel signalling axis transducing metabolic disruption to structural collapse, with consequent impairment of cristae-dependent metabolic enzymes, and provides a potential molecular target for HF intervention.

## Results

### OXPHOS Dysfunction Triggers PHB1/2 Scaffold Disassembly and Cristae Disruption in Heart Failure

Mitochondrial dysfunction, manifesting as reduced OXPHOS function and the structural disruption of cristae, represents a core feature of HF^6^. However, the molecular mechanisms by which OXPHOS dysfunction actively triggers cristae remodelling remain unclear. To investigate this question, we first performed transmission electron microscopy (TEM) analysis on myocardial samples from patients with heart failure with reduced ejection fraction (HF). Compared with non-failing controls (NHF), the cardiomyocyte mitochondria of HF patients exhibited marked structural disorganisation in the cristae, characterised by reduced cristae density and a reduced mitochondrial area (Fig. 1a–b, and Supplementary Table 1). Concurrently, in a transverse aortic constriction (TAC)-induced mouse model of HF, OROBOROS high-resolution respirometry confirmed significantly diminished cardiac mitochondrial oxidative phosphorylation capacity (Extended Data Fig. 1a-b).

**Figure 1.**
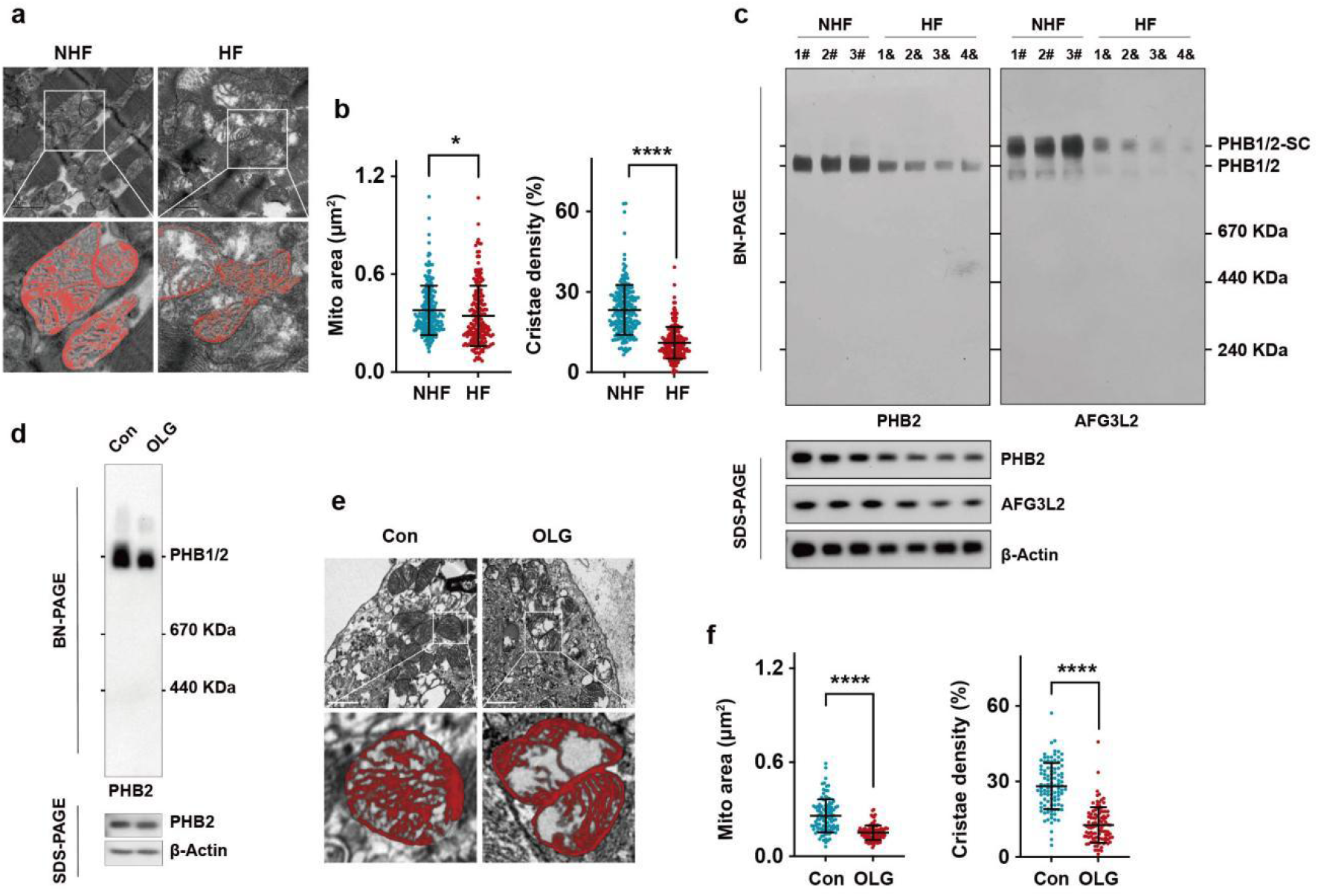
|. OXPHOS Dysfunction Triggers PHB1/2 Scaffold Disassembly and Cristae Disruption in Heart Failure. **a,** Representative transmission electron microscopy (TEM) images of cardiomyocyte mitochondria from non-heart failure (NHF) and heart failure (HF) patients with reduced ejection fraction. **b,** Quantification of mitochondrial area (μm², left) and cristae density (%, right) is shown. Scale bar, 1 μm; n = 100 mitochondria per group (NHF: 3 patients, 5 images per patient, 6–7 mitochondria per image; HF: 4 patients, 5 images per patient, 5 mitochondria per image). **c,** Blue native polyacrylamide gel electrophoresis (BN-PAGE) analysis of PHB1/2 and PHB1/2 super-complex (PHB1/2-SC) assembly in cardiac tissues from NHF and HF patients. **d,** BN-PAGE analysis of PHB1/2 assembly in neonatal mouse cardiomyocytes (NMCMs) treated with vehicle control (Con) or oligomycin (OLG). **e-f,** Representative TEM images (**e**) and the quantification (**f**) of mitochondrial ultrastructure in NMCMs treated with vehicle control or OLG. Scale bar, 0.5 μm; n = 100 mitochondria per group (5 images × 5 mitochondria per image, 4 mice per group). Data are presented as mean ± SD from one representative experiment of at least three independent experiments. Statistical significance was determined by two-tailed unpaired Student’s t-tests (**b**, **f**). *P < 0.05; ****P < 0.0001.

The structural maintenance of cristae depends on the coordinated action of multiple protein complexes; of these, the PHB1/2 (Prohibitin 1/2) complex serves as an inner membrane scaffold protein critical for the morphological stability of cristae^9^. Blue native polyacrylamide gel electrophoresis (BN-PAGE) revealed markedly reduced PHB1/2 super-complex (PHB1/2-SC) assembly in failing human hearts (Fig. 1c). This finding suggests that disassembly of the PHB1/2 complex may represent a key link connecting OXPHOS dysfunction to the structural disruption of cristae.

To verify whether OXPHOS dysfunction can directly cause PHB1/2 disassembly and cristae disruption, we treated neonatal mouse cardiomyocytes (NMCMs) with oligomycin (OLG) to inhibit ATP synthase. Following OLG treatment, BN-PAGE showed significantly reduced PHB1/2-SC assembly (Fig. 1d), while TEM revealed obvious structural disorganisation in the cristae (Fig. 1e-f). Consistent results were observed in the AC16 human cardiomyocyte cell line (Extended Data Fig. 1c-f). Collectively, these data demonstrate that OXPHOS dysfunction can directly trigger disassembly of the PHB1/2 complex and structural disruption in cristae, although the specific molecular mechanisms mediating this process remain unclear.

### NDUFA13-K7 Acetylation Mediates the Coupling Between OXPHOS Dysfunction and PHB1/2 Disassembly

To identify the molecules that mediate signal transmission between OXPHOS dysfunction and PHB1/2 disassembly, we employed immunoprecipitation-mass spectrometry (IP-MS) to compare PHB2 interacting protein profiles before and after OLG treatment. Volcano plot analysis revealed several proteins with significantly enhanced PHB2 interaction after OLG treatment; of these, the complex I accessory subunit NDUFA13 was of particular interest as it belongs to the rate-limiting OXPHOS complex I, and no prior reports had implicated Complex I subunits in direct regulation of cristae scaffold proteins such as prohibitin (Fig. 2a-b). Co-IP experiments with neonatal mouse cardiomyocytes confirmed that multiple OXPHOS inhibition modalities that reduced the NAD^+^/NADH ratio led to enhanced NDUFA13-PHB2 interaction, including H_2_O_2_, rotenone (a complex I inhibitor), antimycin A (a complex III inhibitor), oligomycin (a complex V inhibitor), and sodium azide (a complex IV inhibitor) (Fig. 2c), thus suggesting that this interaction represents a universal response to respiratory dysfunction.

**Figure 2.**
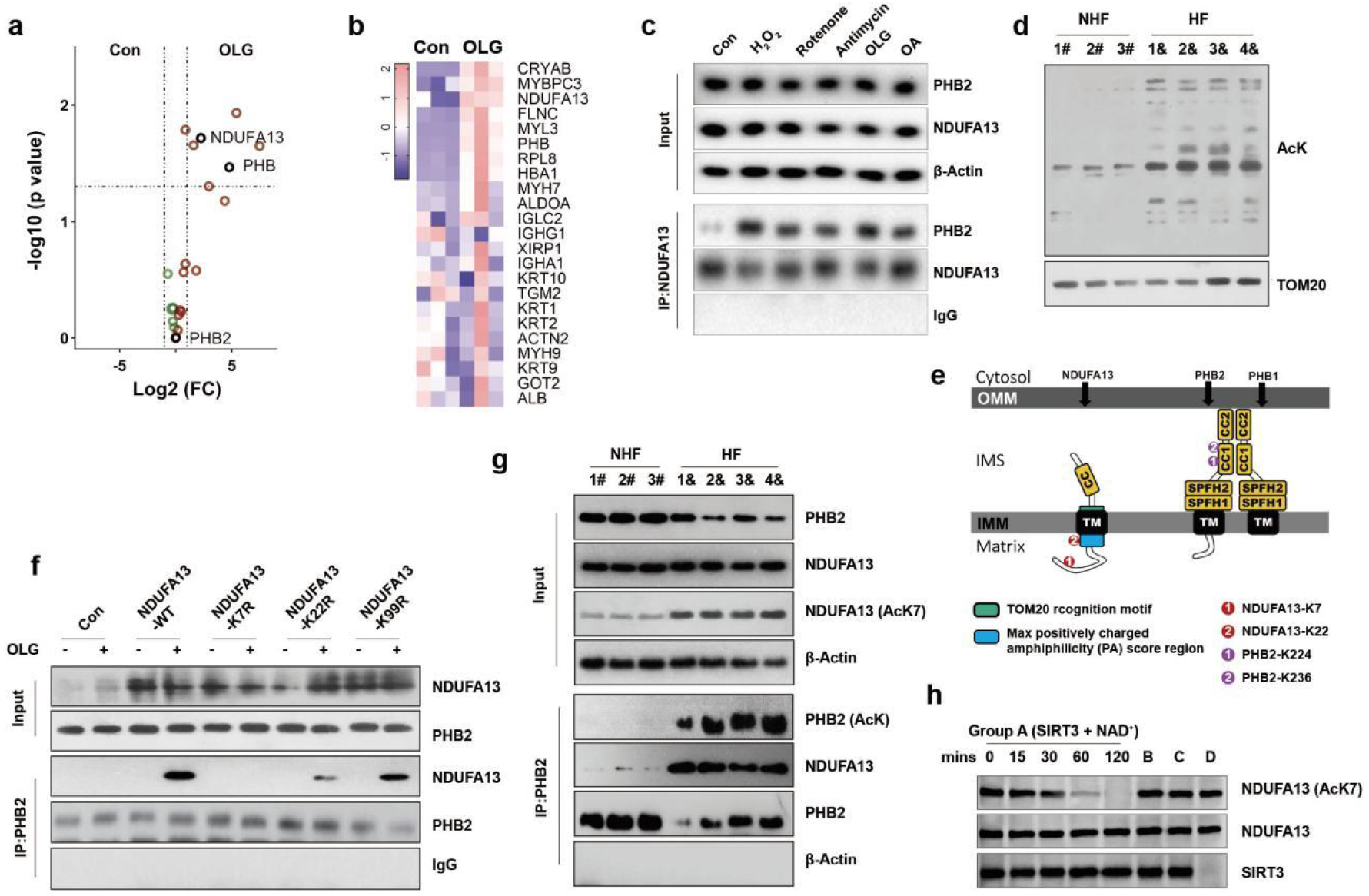
|. NDUFA13-K7 Acetylation Couples OXPHOS Dysfunction to PHB1/2 Disassembly via SIRT3 Inactivation. **a-b,** Interactome analysis following endogenous PHB2 immunoprecipitation in OLG-treated AC16 cells. Differential protein interactions are shown as a volcano plot (**a**) and heatmap (**b**). **c,** Co-immunoprecipitation (Co-IP) analysis of NDUFA13-PHB2 interaction in NMCMs treated with indicated OXPHOS inhibitors: H_2_O_2_, rotenone (Rot), antimycin A (AA), oligomycin (OLG), and sodium azide (NaN_3_). **d,** Western blot analysis of global mitochondrial protein acetylation levels in isolated cardiac mitochondria from NHF and HF patients. **e,** Mass spectrometry quantification of the acetyl-lysine levels on NDUFA13 and PHB2 in vehicle- and OLG-treated AC16 cells. Schematic illustrations depict the spatial localisation and structural features of acetylation site-containing domains. **f,** Co-IP analysis of the interaction between Flag-tagged NDUFA13 variants (WT, K7R, K22R, and K99R) and endogenous PHB2 in NDUFA13-deficient MEF cells. **g,** Western blot and Co-IP analysis of NDUFA13-K7 acetylation, PHB2 acetylation, and NDUFA13-PHB2 interaction in cardiac tissues from healthy controls and HF patients. **h,** *In vitro* deacetylation of endogenous NDUFA13-K7 by recombinant SIRT3. Endogenous NDUFA13 was immunoprecipitated from OLG-treated NMCMs and incubated with recombinant SIRT3 + NAD⁺ (Group A, time course 0–120 min) or negative controls: heat-inactivated SIRT3 (B), no NAD⁺ (C), or no SIRT3 (D). Immunoblots show K7 acetylation (AcK7, upper) and total NDUFA13 (lower). Data shown are from one representative experiment with at least three replicates.

A major metabolic consequence of OXPHOS dysfunction is a reduction in NAD^+^/NADH ratio, which leads to the reduced activity of deacetylases such as SIRT3 and the subsequent hyperacetylation of mitochondrial proteins^6,14^. Western blot analysis further confirmed that global mitochondrial protein acetylation levels were significantly elevated in failing hearts (Fig. 2d). Based on this, we hypothesised that protein acetylation may regulate the interaction between NDUFA13 and PHB2. Mass spectrometry identified lysine 7 (K7) and lysine 22 (K22) of NDUFA13, and lysine 224 (K224) and lysine 236 (K236) of PHB2, as major acetylation sites (Fig. 2e, and Extended Data Fig. 2a-b). To directly verify the function of each acetylation site in regulating interaction, we next constructed several deacetylation-mimetic mutants: NDUFA13-K7R, NDUFA13-K22R, and NDUFA13-K99R (negative control). Co-IP experiments demonstrated that K7R almost completely blocked the NDUFA13-PHB2 interaction, while K22R and K99R retained the interaction (Fig. 2f). Further expression of the PHB2-WT or PHB2-K224R/K236R double mutants in HEK293T-PHB2^-/-^ cells revealed that the double mutation reduced but still preserved NDUFA13-PHB2 interaction (Extended Data Fig. 2c), thus indicating that PHB2 K224/K236 acetylation provided auxiliary enhancement while NDUFA13-K7 was the dominant site.

Using a K7 acetylation-specific antibody (Extended Data Fig. 2d), we performed Co-IP experiments with cardiac samples from healthy individuals and HF patients and found that the levels of K7-acetylated NDUFA13 were significantly increased in the HF group. Following immunoprecipitation with a PHB2 antibody, the levels of acetylated PHB2 were markedly elevated in the HF group, as were the levels of bound NDUFA13 (Fig. 2g). Consistently, site-specific IP/WB in TAC-treated mice confirmed a significant elevation of NDUFA13-K7 acetylation in failing hearts (Extended Data Fig. 3a), with successful model induction verified by echocardiography (Extended Data Fig. 3b-d). To directly validate the upstream deacetylase, we performed an *in vitro* deacetylation assay in which endogenous NDUFA13, immunoprecipitated from OLG-treated NMCMs, was incubated with recombinant SIRT3 and NAD⁺. Immunoblotting with the anti-acetyl-NDUFA13-K7 antibody revealed a progressive, time-dependent reduction in K7 acetylation (Fig. 2h). SIRT3 deacetylase activity was markedly reduced in pressure-overloaded cardiac tissue in vivo (Extended Data Fig. 3e), indicating that NAD⁺ depletion-driven SIRT3 inactivation is the physiological trigger for K7 hyperacetylation in the failing heart. Collectively, these results demonstrate that NDUFA13-K7 and PHB2 acetylation are elevated in HF, thereby enhancing their interaction strength, and establish NDUFA13-K7 as the dominant acetylation site governing this process.

### K7 Acetylation Induced Conformational Changes in NDUFA13, Intermembrane Space Translocation, and Dynamic Dissociation from Complex I

The K7 site of NDUFA13 is located in its matrix-facing N-terminal exposed region. Structural analysis of Complex I (PDB: 6G2J)^18^ showed that this site is proximal to the NADH binding site, positioned ideally for sensing redox status (Extended Data Fig. 4a). To investigate how K7 acetylation affects protein conformation, we next performed 200 ns molecular dynamics simulations. Analysis revealed that when compared to the non-acetylated form, K7 acetylation significantly reduced the radius of gyration (Rg) and solvent accessible surface area (SASA), while increasing intramolecular hydrogen bond number, thus resulting in more compact protein folding (Fig. 3a). Structural visualisation and residue–residue interaction energy matrix analysis further revealed marked conformational differences between non-acetylated and acetylated NDUFA13 states (Fig. 3b-c).

**Figure 3.**
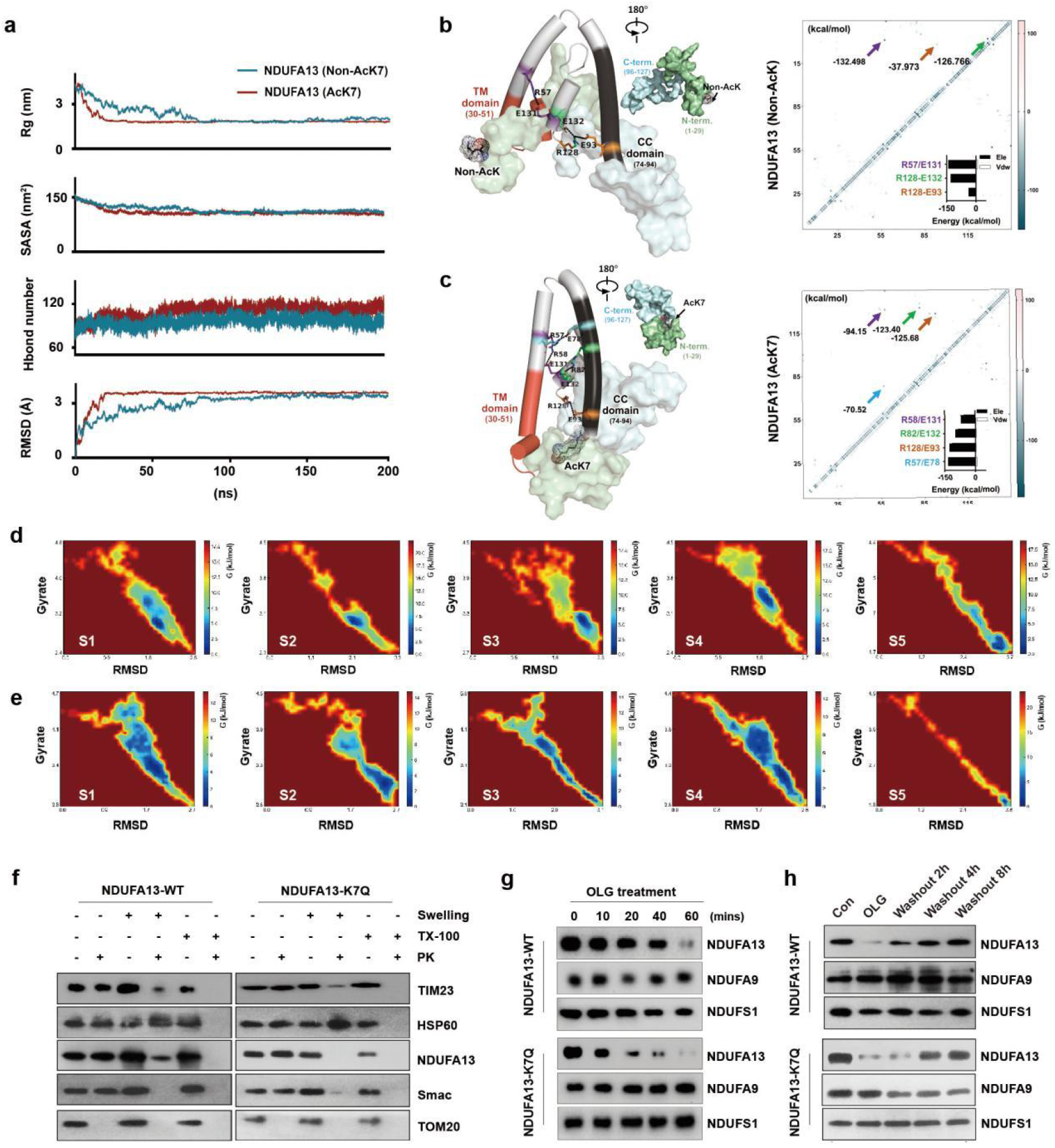
|. K7 Acetylation induced NDUFA13 conformational compaction, intermembrane space translocation, and dynamic dissociation from complex I. **a,** Molecular dynamics simulations (200 ns) comparing folding dynamics of acetylated (AcK7) and non-acetylated (Non-AcK7) NDUFA13. Parameters include radius of gyration (Rg, nm), solvent-accessible surface area (SASA, nm²), intramolecular hydrogen bond numbers, and root-mean-square deviation (RMSD, Å). **b-c,** Structural and energetic analysis of non-acetylated (Non-AcK7, **b**) and acetylated (AcK7, **c**) NDUFA13 based on 200 ns molecular dynamics simulations. Left panels: Three-dimensional structural models depicting transmembrane (TM, dark blue) and coiled-coil (CC, teal) domains as cylindrical helices, with C-terminal (light blue) and N-terminal (orange) regions shown as surface representations. Electrostatic interactions are colour-coded as indicated. Right panels: Residue–residue interaction energy matrices (kcal/mol), with colour gradients representing attractive (red, negative values) and repulsive (blue, positive values) forces. **d-e,** Free energy landscape (FEL) analysis showing global energy minima (ΔG, kJ/mol) of acetylated NDUFA13 (AcK7) (**d**) and non-acetylated NDUFA13 (Non-AcK7) (**e**) as a function of RMSD (nm) and radius of gyration (Rg, nm) over 200 ns molecular dynamics simulations. **f,** Submitochondrial localisation analysis of NDUFA13 variants in NDUFA13-deficient MEF cells. Isolated mitochondria were subjected to hypotonic swelling to disrupt the outer mitochondrial membrane (OMM), followed by membrane solubilisation with Triton X-100, with or without proteinase K (PK; 50 μg/mL) treatment. Smac (intermembrane space marker) and Hsp60 (matrix marker) served as compartment controls. **g,** BN-PAGE analysis of complex I assembly dynamics in NDUFA13-deficient MEF cells expressing Flag-tagged NDUFA13 variants (WT, K7Q, and K7R) following OLG treatment (12 h) and subsequent washout at indicated time points. **h,** Quantitative analysis of NDUFA13-K7Q incorporation rate into complex I at indicated time points following transient transfection into NDUFA13-deficient MEF cells. Data shown are from one representative experiment with at least three replicates.

Free energy landscape (FEL) analysis provided a thermodynamic perspective on the conformational effects of K7 acetylation. Non-acetylated NDUFA13 exhibited a relatively flat energy landscape, suggesting high conformational flexibility while K7 acetylation produced a clear energy well in the landscape, thus indicating stabilisation of the protein in a specific compact conformation (Fig. 3d-e). Lowest energy conformation extraction, structural snapshots every 20 ns during simulation, and DSSP secondary structure time-course analysis further confirmed that K7 acetylation significantly enhanced the stability of the NDUFA13 protein (Extended Data Fig. 4b-g).

Submitochondrial fractionation experiments further revealed the effect of K7 acetylation on NDUFA13 localisation. First, immunofluorescence analysis revealed that K7 acetylation did not cause the escape of NDUFA13 to the cytoplasm. Even under conditions of OXPHOS inhibition, both NDUFA13-K7Q (acetylation-mimetic mutant) and NDUFA13-WT remained localised to the mitochondria (Extended Data Fig. 5a). Subsequently, the combination of submitochondrial fractionation and proteinase K protection assays revealed that NDUFA13-K7Q was more susceptible to degradation than NDUFA13-WT, thus suggesting its preferential exposure in the intermembrane space (IMS) (Fig. 3f). In alkaline extraction experiments, NDUFA13-WT was primarily retained in the pellet, as with the inner membrane integral protein TIM23; NDUFA13-K7Q exhibited dual localisation, appearing in both the pellet and supernatant (Extended Data Fig. 5b).

Dynamic analysis further revealed that OLG treatment significantly accelerated NDUFA13-K7Q dissociation from complex I (Fig. 3g); following OLG washout or under conditions of exogenous expression, the reassembly rate was markedly slower than the wild-type (Fig. 3h, and Extended Data Fig. 5c). Despite altered kinetic behaviour, the final assembly levels of complex I and enzymatic activity in cells expressing K7Q and K7R mutants did not differ significantly from the wild-type (Extended Data Fig. 5d-e). These results indicated that K7 acetylation does not disrupt the catalytic function of complex I, but rather converted NDUFA13 into a highly OXPHOS status-sensitive signalling molecule by modulating its dynamic residence time within complex I, thus enabling the rapid translocation and execution of PHB1/2 scaffold remodelling during dysfunction.

Collectively, these results demonstrated that K7 acetylation induced NDUFA13 conformational compaction and partial translocation to the IMS, thereby altering its localisation and accessibility in the membrane environment, thus providing a structural basis for subsequent aberrant binding to PHB2.

### Acetylated NDUFA13 Competitively Disrupted PHB1/2 Assembly Through Coiled-Coil Domains

According to recently reported cryo-ET structures, PHB1 and PHB2 form heterodimers through parallel coiled-coil (CC) domain interactions, representing the fundamental unit of ring complex assembly^11^. As shown in Extended Data Fig. 2b, OXPHOS dysfunction led to elevated acetylation levels at the PHB2 K224 and K236 sites. Structural analysis of the PHB1/2 complex further revealed that K224 and K236 were located in the CC domain of PHB2; as charged amino acids, these participated in forming the electrostatic network between PHB1-PHB2 (Extended Data Fig. 6a). Acetylation of PHB2-K224/K236 disrupted this electrostatic network, leading to disassembly of the PHB1/2 complex (Fig. 4a).

**Figure 4.**
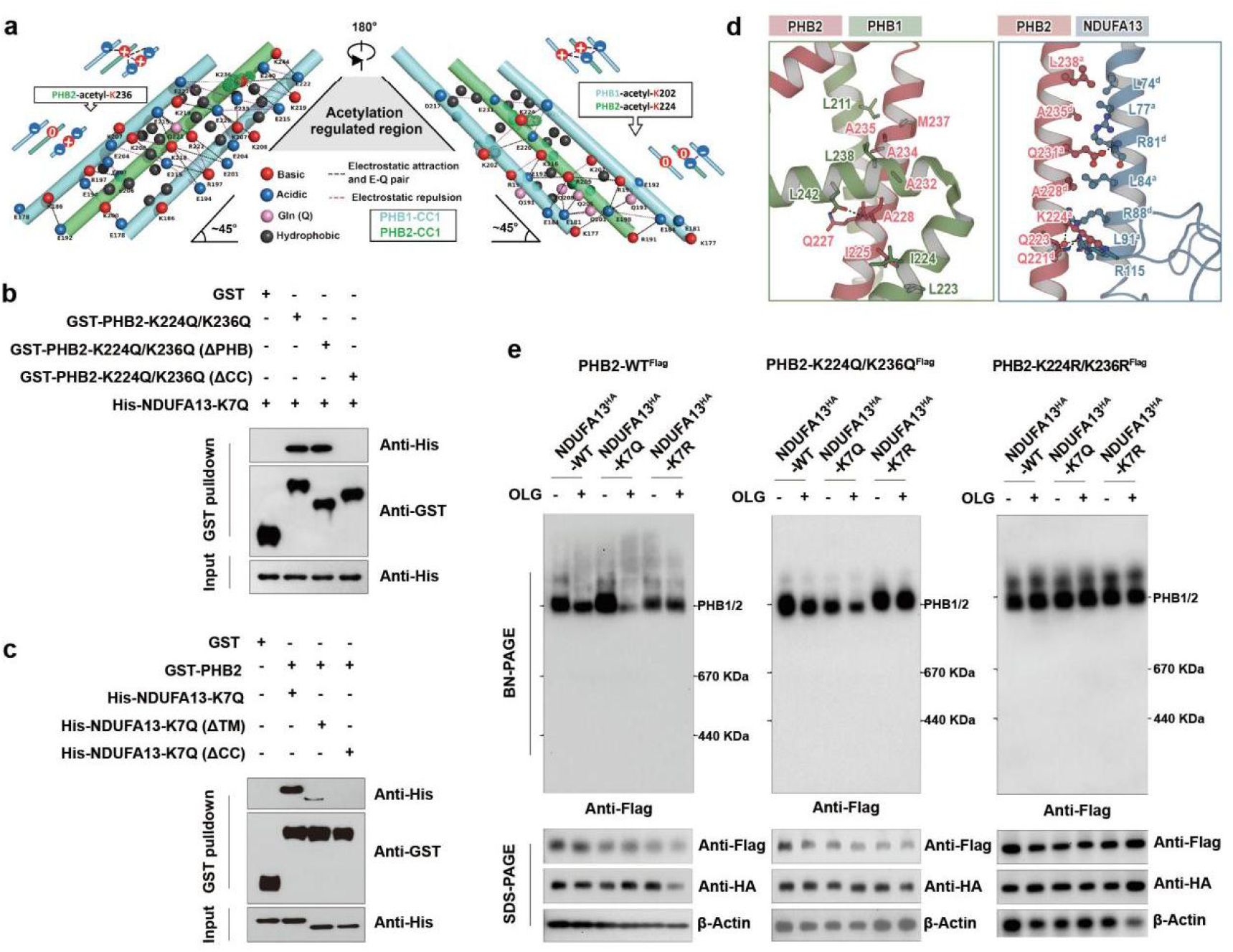
|. Acetylated NDUFA13 competitively disrupted PHB1/2 assembly through coiled-coil domain interactions. **a,** Schematic representation of electrostatic network interactions between the coiled-coil (CC1) domains of PHB1 and PHB2, illustrating alterations in interaction forces upon acetylation at PHB1-K202 and PHB2-K224/K236. **b,** GST pulldown analysis of His-NDUFA13 (WT or K7Q) binding to GST-PHB2 variants, including wild-type, PHB domain deletion (ΔPHB), and coiled-coil domain deletion (ΔCC). **c,** GST pulldown analysis of GST-PHB2 (WT or K224Q/K236Q) binding to His-NDUFA13 variants, including wild-type, transmembrane domain deletion (ΔTM), and coiled-coil domain deletion (ΔCC). **d,** Structural comparison of PHB1-PHB2 and NDUFA13-PHB2 coiled-coil interfaces. Left: Magnified view of native interactions between PHB2 (pink) and PHB1 (green) based on cryo-ET structure (PDB: 9O6S). Right: Magnified view of predicted interactions between PHB2 (pink) and NDUFA13 (blue) from molecular docking analysis. Hydrophobic residues at heptad repeat positions “a” and “d” are shown in ball-and-stick representation and labelled with superscript notations. **e,** BN-PAGE analysis of PHB1/2 complex and PHB1/2 super-complex (PHB1/2-SC) assembly in NDUFA13-deficient MEF cells co-expressing indicated PHB2 (WT or K224R/K236R) and NDUFA13 (WT, K7Q, or K7R) variants, with or without OLG treatment. Blots shown are from one representative experiment with at least three replicates.

Next, we investigated the molecular basis underlying NDUFA13-PHB2 interaction. Sequence and structural analysis revealed that NDUFA13 also possessed a conserved coiled-coil (CC) domain, with the “a” and “d” positions enriched in hydrophobic residues, thus conforming to the classical Knobs-into-Holes (KIH) packing mode and suggesting potential coiled-coil-mediated competitive binding to PHB2 (Extended Data Fig. 6b)^19,20^. To identify the domains mediating interaction, we next constructed a series of deletion mutants and performed GST pulldown experiments. Analysis revealed that deletion of either the PHB2 CC region or the NDUFA13 CC region completely abolished interaction between the two proteins, while deletion of other domains had no effect (Fig. 4b-c). This indicated that CC domains are both necessary and sufficient for NDUFA13-PHB2 interaction.

Molecular docking simulations were then performed to screen six potential CC–CC docking modes (Extended Data Fig. 6c). Binding energy analysis indicated that the antiparallel-1 configuration exhibited the most stable binding (Extended Data Fig. 6d). Structural comparison of the NDUFA13-PHB2 interface with the native PHB1-PHB2 interface revealed significant overlap between the two structures (Fig. 4d), implying that NDUFA13 competed with PHB1 for the same binding interface on PHB2; when K7-acetylated NDUFA13 bound to PHB2, it displaced PHB1 from the complex, leading to disassembly of the PHB1/2 complex. This finding was consistent with our earlier observation that PHB2-K224R/K236R mutation (preventing acetylation at these two sites) reduced NDUFA13-PHB2 formation under OLG treatment (Extended Data Fig. 2c). Furthermore, blue native polyacrylamide gel electrophoresis (BN-PAGE) following Flag-tagged protein elution demonstrated that the inhibition of acetylation at either NDUFA13-K7 or PHB2-K224R/K236R rescued disruption of the PHB1/2 complex under OLG treatment (Fig. 4e). In summary, NDUFA13 formed competitive interactions with the CC region of PHB2 through its own CC domain; under conditions in which K7 acetylation provided primary initiation and PHB2 acetylation provided auxiliary enhancement, NDUFA13 effectively displaced PHB1, leading to disassembly of the PHB1/2 complex.

### NDUFA13-K7 Acetylation was Sufficient to Induce the Remodelling of Cristae and Accelerate the Progression of HF

To verify whether NDUFA13-K7 acetylation was sufficient to drive downstream phenotypes of PHB1/2 disassembly, we next reconstituted wild-type (WT), acetylation-mimetic (K7Q), and deacetylation-mimetic (K7R) NDUFA13 variants in NDUFA13-deficient MEF cells. TEM revealed that MEF cells expressing NDUFA13-K7Q exhibited reduced mitochondrial cristae density compared with NDUFA13-WT-expressing cells following OLG treatment, while NDUFA13-K7R expression protected cells from structural disruption of the cristae (Fig. 5a–b). Consistently, MitoSOX fluorescence showed increased mitochondrial ROS production in the K7Q group and rescue in the K7R group following OLG treatment (Extended Data Fig. 7b). These results demonstrated that NDUFA13-K7 acetylation alone was sufficient to induce pathological cristae remodelling, oxidative stress, and mitochondrial dysfunction.

**Figure 5.**
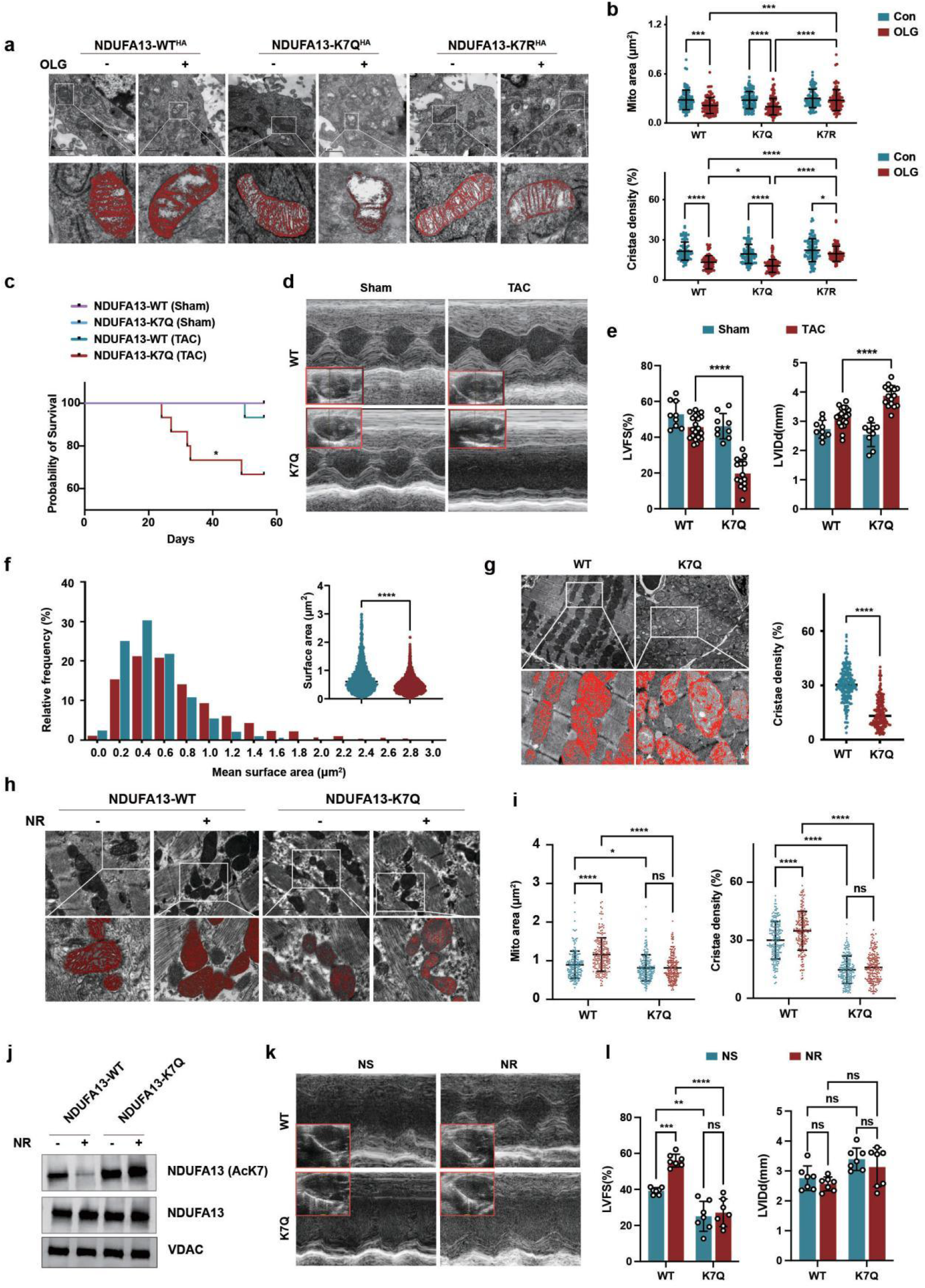
|. NDUFA13-K7 acetylation was sufficient to induce cristae remodelling and accelerate the progression of heart failure. **a,** Representative TEM images of mitochondrial ultrastructure in NDUFA13-deficient MEF cells reconstituted with wild-type (WT), K7Q, or K7R NDUFA13, with or without OLG treatment. **b**. Quantification of mitochondrial area (μm², upper) and cristae density (%, lower) is shown. Scale bar, 1 μm; n = 100 mitochondria per group (5 images × 5 mitochondria per image, 4 mice per group). **c,** Kaplan-Meier survival analysis of cardiomyocyte-specific NDUFA13-K7Q knock-in mice (Myh6^Cre+^; NDUFA13-K7Q^fl/fl^, n = 18) and control littermates (Myh6^Cre−^; NDUFA13-K7Q^fl/fl^, n = 12) following sham or TAC surgery. **d,** Representative M-mode echocardiographic images of the left ventricle from indicated experimental groups. **e,** Quantification of left ventricular fractional shortening (LVFS, left) and left ventricular internal diameter at end-diastole (LVIDd, right) is shown. n = 8–10 mice per group. **f,** Frequency distribution analysis of mitochondrial cross-sectional areas from cardiomyocytes of NDUFA13-K7Q knock-in and control mice following TAC surgery, with quantification of mean mitochondrial area.n > 2000 mitochondria per group (25 images × 80∼ mitochondria per image, 4 mice per group). **g,** Representative TEM images of cardiac mitochondria from indicated experimental groups, with corresponding quantification of cristae density. Scale bar, 2 μm; n = 200 mitochondria per group (10 images × 5 mitochondria per image, 4 mice per group) for cristae density quantification. **h,** Representative TEM images of mitochondrial ultrastructure in cardiac tissues from NDUFA13-K7Q knock-in and control mice following TAC surgery treated with or without NR. **i,** Quantification of mitochondrial area (μm², left) and cristae density (%, right) is shown. Scale bar, 0.5 μm; n = 200 mitochondria per group (10 images × 5 mitochondria per image, 4 mice per group) for cristae density quantification. **j,** IP/WB showing AcK7 (upper), total NDUFA13 (middle), and VDAC loading control (lower). **k,** Representative M-mode echocardiographic images of the left ventricle from indicated experimental groups. **l,** Quantification of left ventricular fractional shortening (LVFS, left) and left ventricular internal diameter at end-diastole (LVIDd, right) is shown. n = 7 mice per group. Data are presented as mean ± SD from one representative experiment of at least three independent experiments. Statistical significance was determined by two-way ANOVA with multiple comparison (**b**, **e**, **i, l**), log-rank test (**c**), or two-tailed unpaired Student’s t-test (**f**, **g**). *P < 0.05; ***P < 0.001;****P < 0.0001; ns, not significant.

To investigate the pathophysiological significance of this mechanism *in vivo*, we generated cardiomyocyte-specific NDUFA13-K7Q knock-in mice (Myh6^Cre+^; NDUFA13-K7Q^flox/flox^) (Extended Data Fig. 7a). Compared to control mice (Myh6^Cre-^; NDUFA13-K7Q^flox/flox^), K7Q mice exhibited significantly reduced survival following TAC surgery (Fig. 5c). Histological evaluation revealed marked adverse cardiac remodelling in TAC-treated K7Q knock-in mice, including ventricular dilation, increased fibrosis, and cardiomyocyte hypertrophy (Extended Data Fig. 7c). Echocardiography demonstrated severely impaired cardiac function in K7Q knock-in mice following TAC, manifesting as significantly reduced LVFS and increased LVIDd (Fig. 5d–e). TEM and mitochondrial cross-sectional area frequency distribution analysis further confirmed significantly reduced mitochondrial area and cristae density in TAC-treated K7Q knock-in mice compared with controls (Fig. 5f–g).

To determine whether the cardioprotective effect of NAD⁺ repletion requires NDUFA13-K7 deacetylation, WT and NDUFA13-K7Q knock-in mice underwent TAC and received NR or normal saline (NS) daily for 4 weeks. In WT+TAC mice, NR treatment significantly reduced NDUFA13-K7 acetylation, restored PHB1/2 assembly, improved cardiac function, and recovered cristae density relative to NS-treated controls (Fig. 5h–l, and Extended Data Fig. 7d). In contrast, none of these parameters were rescued by NR in K7Q knock-in mice, which remained indistinguishable from K7Q+TAC+NS controls across all endpoints. As the K7Q substitution constitutively mimics the acetylated state and is refractory to SIRT3-mediated deacetylation, these findings demonstrate that the cardioprotective effect of NR is strictly dependent on NDUFA13-K7 deacetylation.

Collectively, these *in vitro* and *in vivo* findings demonstrated that NDUFA13-K7 acetylation significantly accelerated pressure overload-induced HF progression by exacerbating cristae structural collapse through disruption of the prohibitin scaffold. Furthermore, the failure of NR to rescue the K7Q phenotype conclusively establishes NDUFA13-K7 deacetylation as an essential downstream effector of NAD⁺-mediated cardioprotection, mechanistically distinguishing this axis from non-specific pleiotropic effects of NAD⁺ repletion.

### NDUFA13-K7Q Mutation Disrupts Tricarboxylic Acid (TCA) Cycle and Fatty Acid Transport in Cardiomyocytes

To determine whether NDUFA13-K7 acetylation-induced cristae remodelling translated into functional metabolic deficits, we performed targeted metabolomics profiling on cardiac tissues from NDUFA13-K7Q knock-in (Myh6^Cre+^; NDUFA13-K7Q^flox/flox^) and control (Myh6^Cre-^; NDUFA13-K7Q^flox/flox^) mice following TAC surgery. This analysis identified 35 metabolites with statistically significant alterations between genotypes (Fig. 6a, Extended Data Fig. 8, and Supplementary Table 2).

**Figure 6.**
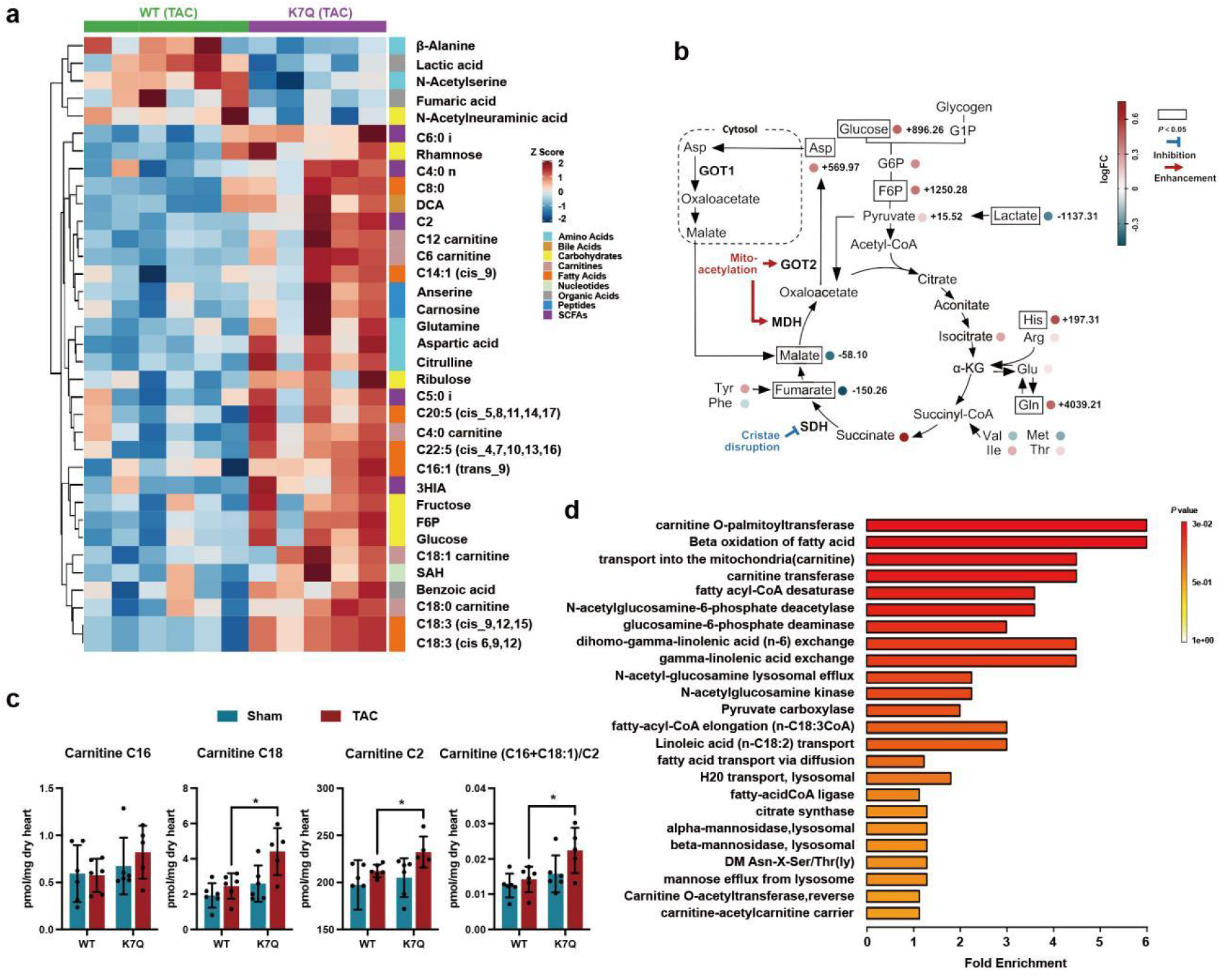
|. NDUFA13-K7Q mutation disrupts TCA cycle and fatty acid metabolism in cardiomyocytes. **a,** Heatmap showing differential metabolite profiles between cardiomyocyte-specific NDUFA13-K7Q knock-in mice (Myh6^Cre+^; NDUFA13-K7Q^fl/fl^) and control littermates (Myh6^Cre−^; NDUFA13-K7Q^fl/fl^) following TAC surgery. **b,** Comparative analysis of glycolytic and TCA cycle intermediates in cardiac tissues of NDUFA13-WT and NDUFA13-K7Q mice post-TAC treatment. Included are the malate-aspartate shuttle and amino acids related to anaplerotic and cataplerotic pathways of the TCA cycle, with differences and fold changes presented numerically and as a heatmap, respectively. **c,** Quantification of C16- and C18:1-carnitines, and the (C16+C18:1)/C2 ratio. **d,** Pathway enrichment analysis bar chart comparing NDUFA13-K7Q and NDUFA13-WT, based on the Predicted Metabolite Sets database. The x-axis bar length denotes fold enrichment, and the bar color indicates the P-value, with red intensity directly correlating to pathway significance. Data are presented as mean ± SD. Statistical significance was determined by two-tailed unpaired Student’s t-test (**c**). *P < 0.05.

TCA cycle intermediates displayed impaired flux in NDUFA13-K7Q hearts, with elevated succinate and diminished fumarate. As succinate dehydrogenase (SDH) is embedded in the cristae membrane and requires intact cristae architecture for optimal activity^5^, this metabolite signature provides direct evidence for functional impairment of SDH-catalysed succinate oxidation, consistent with K7Q-driven cristae disruption. Additionally, significant accumulation of glycolytic intermediates (glucose, G6P, F6P, and pyruvate) indicated enhanced glycolytic flux, consistent with the Warburg-like metabolic shift characteristic of failing hearts (Fig. 6b).

Fatty acid oxidation was profoundly impaired in NDUFA13-K7Q hearts. Long-chain acylcarnitines markedly accumulated, with significantly elevated carnitine-C16 and carnitine-C18:1 levels and an increased (C16+C18:1)/C2 ratio (Fig. 6c), a hallmark of carnitine palmitoyltransferase II (CPT II) dysfunction. Given that CPT II is associated with the cristae membrane and requires intact cristae architecture for optimal activity^21^, this pattern provided direct evidence of defective mitochondrial fatty acid import secondary to cristae disruption. Comprehensive profiling further revealed broad elevations in acylcarnitines and free fatty acids across multiple chain lengths (Extended Data Fig. 9a-c). Z-score analysis provided an integrated visualisation of the global metabolic perturbations in K7Q hearts (Extended Data Fig. 9d). Pathway enrichment analysis using the Small Molecule Pathway Database (SMPDB) revealed significant enrichment of Warburg effect, amino acid metabolism, and TCA cycle pathways (Extended Data Fig. 9e). Further analysis using the Predicted Metabolite Sets library identified pronounced disruption of carnitine transport and carnitine transferase pathways (Fig. 6d), consistent with impaired mitochondrial fatty acid import.

To determine whether NDUFA13-K7Q-driven cristae remodelling was accompanied by corresponding disruption of the inner membrane lipid environment, we performed untargeted cardiolipin (CL) lipidomics by LC-MS/MS on cardiac tissues from all four experimental groups (WT+Sham, WT+TAC, K7Q+Sham, K7Q+TAC). The predominant cardiac CL species tetralinoleoyl-cardiolipin (L4CL; CL 18:2/18:2/18:2/18:2) was significantly reduced in WT+TAC hearts compared with WT+Sham controls, consistent with OXPHOS-driven CL degradation under pressure overload (Extended Data Fig. 10a). Correspondingly, the MLCL/CL ratio, which rises when tafazzin-mediated reacylation is impaired, was progressively elevated across groups, with the highest levels observed in K7Q+TAC hearts, indicating a graded impairment of the CL remodelling cycle that parallels the severity of cristae disruption (Extended Data Fig. 10b). Heatmap analysis of all significantly altered CL species identified 18 CL species that were substantially decreased in K7Q+TAC hearts relative to WT+TAC, with polyunsaturated linoleoyl-containing species showing the greatest depletion (Extended Data Fig. 10c, and Supplementary Table 3). To investigate the enzymatic basis for these findings, we assessed protein levels of tafazzin (TAZ) and cardiolipin synthase (CLS) by Western blot. Both TAZ and CLS were significantly reduced in WT+TAC hearts relative to sham controls, and were further decreased in K7Q+TAC hearts (Extended Data Fig. 10d). Given that TAZ functions within a DNAJC19–TAZ–prohibitin complex in which the PHB scaffold is required for TAZ stabilisation, PHB1/2 disassembly driven by K7 acetylation provides a direct mechanistic explanation for the impaired CL reacylation observed in K7Q+TAC hearts.

## Discussion

The collapse of mitochondrial cristae in HF has long been viewed as a passive consequence of metabolic energy dysfunction, yet the molecular mechanisms underlying this process have yet to be fully elucidated^22,23^. In this study, we identified a novel and active signalling pathway in that the complex I accessory subunit NDUFA13 exerts functionality not merely as a structural component of the electron transport chain, but as a dual-role protein with metabolic sensing capabilities^24^. Our analyses showed that following OXPHOS impairment, the NDUFA13-K7 site underwent hyperacetylation, triggering conformational compaction and translocation into the IMS, subsequently competitively displacing PHB1 through coiled-coil domain interactions to dismantle prohibitin scaffolds. This mechanism directly couples dynamic changes in metabolic status to the structural homeostasis of cristae, revealing a central role for the regulation of acetylation in the pathogenesis of HF.

Our work identified NDUFA13-K7 as a highly sensitive redox rheostat. This site is proximal to the NADH binding pocket and ideally positioned to sense a reduction in the NAD^+^/NADH ratio during HF^25^. We found that the resulting acetylation induced more compact protein conformation and partial translocation to the IMS, significantly altering its dynamic residence behaviour within complex I, accelerating dissociation under stress and delaying reassembly during recovery but without affecting final complex I catalytic activity. This kinetic reprogramming transformed what was originally a stable accessory subunit into a signal transduction molecule, rapidly engaging PHB2 when OXPHOS was impaired.

The competitive nature of this interaction is particularly noteworthy. Both NDUFA13 and PHB1 bind PHB2 through conserved coiled-coil regions, although our computational docking and MM/PBSA analysis suggested that acetylated NDUFA13 adopts an antiparallel orientation with a more favourable binding free energy for PHB2 than PHB1, which could enable displacement of PHB1 from its native parallel heterodimer. Although PHB2 acetylation at K224/K236 provided auxiliary enhancement, the dominant role of NDUFA13-K7 was unequivocal in that K7R deacetylation-mimetic mutation almost completely blocked binding and achieved functional rescue. This finding is highly consistent with recently reported structures of the PHB1/2 complex, supporting the structural feasibility of competitive displacement at the CC1 interface^11^.

We also found that expression of the acetylation-mimetic NDUFA13-K7Q alone was sufficient to induce cristae rarefaction and mitochondrial ROS accumulation *in vitro*, while cardiomyocyte-specific K7Q knock-in mice showed accelerated mortality, adverse remodelling, and severe cristae disruption under pressure overload. These phenotypes are highly consistent with human HFrEF, validating the clinical relevance of this signalling axis^1,26^.

Importantly, our metabolomic analyses provided direct functional evidence that NDUFA13-K7Q-driven cristae remodelling translated into impaired activity of cristae-resident enzymes. The accumulation of succinate with diminished fumarate, and elevated (C16+C18:1)/C2 acylcarnitine ratio with broad FFA accumulation, indicated impaired activity of SDH and CPT II, respectively. SDH, embedded in the inner membrane; and CPT II, associated with the cristae membrane, both require intact cristae architecture for optimal function^27^. The concurrent enrichment of glycolytic intermediates and Warburg effect pathway suggested compensatory metabolic reprogramming characteristic of failing hearts^28^. These findings establish a complete pathophysiological cascade: NDUFA13-K7 acetylation → PHB1/2 disassembly → cristae disruption → impairment of cristae-dependent enzymes → metabolic reprogramming, thereby providing functional validation that structural collapse of cristae directly compromises cardiac energetics.

Our work integrates multiple emerging concepts in the field of cardiac bioenergetics. First, we provide a mechanistic explanation for the long-standing observation of mitochondrial protein hyperacetylation in HF, identifying reversible lysine acetylation as a rapid sensitive switch rather than merely a metabolic byproduct. Second, we revealed a new regulatory layer for prohibitin scaffolds, which were previously viewed primarily as static structural elements or cardiolipin chaperones^9,29^. Third, the dynamic dissociation-reassembly behaviour of NDUFA13 demonstrated how respiratory chain subunits can perform dual functions, serving as both enzymatic components and signalling molecules, a pattern increasingly confirmed for other complex I accessory subunits^30,31^. Fourth, the metabolic consequences we observed, particularly the disruption of fatty acid oxidation and enhanced glycolysis, mirror the well-documented substrate utilisation shift in HF, suggesting that NDUFA13-K7 acetylation may represent an upstream driver of this metabolic inflexibility^32^.

Cardiolipin lipidomics and enzymatic profiling provided further mechanistic insight into this pathway. In addition to their well-characterised structural role in cristae maintenance, prohibitin scaffolds serve as platforms for CL remodelling, coordinating DNAJC19 and tafazzin within a supramolecular complex that facilitates reacylation of MLCL to mature TLCL26. We found that PHB1/2 disassembly in pressure-overloaded K7Q hearts was associated with reduced tafazzin protein abundance and a corresponding MLCL/CL imbalance that was absent under baseline conditions, indicating that the CL remodelling defect depends on concurrent haemodynamic stress. The reduction in cardiolipin synthase in these hearts suggests that de novo CL biosynthesis is also affected, though whether this reflects a direct consequence of cristae disruption or a broader impairment of mitochondrial biogenesis under pressure overload warrants further investigation. These data indicate that NDUFA13-K7 acetylation, by dismantling prohibitin scaffolds, initiates a secondary lipid remodelling defect that compromises the CL composition of the inner membrane and may amplify cristae instability through a feed-forward mechanism.

From a therapeutic perspective, the amenability of this signalling axis to intervention warrants further attention. As a post-translational modification, NDUFA13-K7 acetylation can theoretically be modulated *via* the regulation of upstream enzyme activity. Strategies that enhance NAD^+^ bioavailability (such as nicotinamide riboside supplementation) have shown promise for improving cardiac function in clinical trials of early-phase HF^33–35^, while the development of SIRT3-specific activators is actively progressing in preclinical research^36–38^. These interventions may promote NDUFA13-K7 deacetylation, maintain PHB1/2 assembly, and protect the integrity of cristae. Furthermore, selective interference with the NDUFA13-PHB2 coiled-coil interface represents another potential direction, although achieving specificity remains challenging.

Our study has several limitations. First, the human cohort was small (n = 3 non-failing and 4 failing hearts) with clinically heterogeneous controls, so the human data should be interpreted as supporting clinical correlation rather than proof, with the core mechanism resting on the cell and K7Q knock-in mouse models. Second, we relied mainly on acetylation-mimetic (K7Q) and non-acetylatable (K7R) substitutions rather than endogenous site-specific acetylation. Future studies employing base-editing strategies to introduce K7Q or K7R modifications at the endogenous locus would provide more definitive mechanistic evidence. Additionally, future work should also investigate whether this signalling axis operates in other high-energy-demand tissues and whether it is reversibly engaged during ischaemia-reperfusion injury.

## Conclusion

We identified a novel acetylation-dependent signalling cascade in which NDUFA13-K7 served as an OXPHOS sentinel, competitively hijacking PHB2 during metabolic stress to dismantle prohibitin scaffolds and drive the pathological remodelling of cristae in HF. The resulting cristae disruption impaired the activity of cristae-resident enzymes, leading to TCA cycle dysfunction and defective fatty acid oxidation. This discovery establishes a direct molecular bridge between bioenergetic failure and the structural collapse of the inner mitochondrial membrane, thus providing new mechanistic insights and therapeutic opportunities for a disease that urgently requires novel interventional strategies.

## Acknowledgements

We would like to acknowledge the supporting of Prof. Qiming Sun from Zhejiang university. We thank Dandan Song and Menghan Zhang in the Center of Cryo-Electron Microscopy (CCEM), Zhejiang University for their technical assistance on Transmission Electron Microscopy. We thank Ms. Yingying Huang and Ms. Yanwei Li from the Core Facilities, Zhejiang University School of Medicine for their technical support.

## Author Contributions Statement

J.N., X.H., B.L. and Z.S. conceived, designed and supervised the overall project. J.N., Z.S., Y.L., J.S. and H.C. performed most experiments and analysed data. X.Y., Z.X., J.Y., Y.W., C.K., T.R. and J.J. contributed to in vitro experiments, mouse model generation, and biochemical analyses. D.X. and X.H. performed molecular dynamics simulations and structural analyses, with assistance from X.W. and B.W., X.K., Z-Z.L. and Z.Z. assisted with mitochondrial isolation, respirometry, and imaging studies. D.Z. provided expertise in echocardiography and histological analyses. J.N. and Z.S. wrote the manuscript.

## Data Availability

The untargeted cardiolipin lipidomics data have been deposited to MetaboLights under accession code MTBLS14645, and the targeted metabolomics data under accession code MTBLS1463. Previously published structural coordinates used in this study are available from the Protein Data Bank under accession codes 9O6S (human prohibitin complex) and 6G2J (mouse mitochondrial complex I). Source data for all main and Extended Data figures are provided with this paper. Human cardiac tissue datasets are subject to controlled access to protect patient privacy; de-identified data are available from the corresponding author (J.N.) upon reasonable request, subject to approval by the Ethical Committee for Human Studies of the Second Affiliated Hospital of Zhejiang University School of Medicine and completion of a data transfer agreement. All other data supporting the findings of this study are available within the article and its Supplementary Information.

## Code availability

No custom code or algorithms were used in this study. All data were analysed using publicly available software packages as described in the Methods, including GROMACS (v2018.3), the g_mmpbsa tool (v1.0), the HDOCK server, CHARMM-GUI, MODELLER (v10.4), AmberTools 23, MaxQuant (v2.4.2.0), MS-DIAL, QuanMET (v2.0), MetaboAnalyst 5.0, R (v4.2.0) and GraphPad Prism (v9.5.1).

## Methods

### Reagents

Oligomycin (OLG), rotenone, antimycin A, sodium azide, and hydrogen peroxide (H_2_O_2_) were obtained from Sigma-Aldrich (St. Louis, MO, USA). MitoSOX Red were purchased from Invitrogen (Carlsbad, CA, USA). MitoTracker Deep Red was from Thermo Fisher Scientific. Beta-Nicotinamide Riboside (NR) was purchased from Medchemexpress (MCE, USA).

### Antibodies

The following antibodies were used: rabbit anti-acetyl-lysine (Cell Signaling Technology, #9441, 1:1000), rabbit anti-TOM20 (Cell Signaling Technology, #42406, 1:500), rabbit anti-PHB2 (Abcam, ab71970, 1:1000), rabbit anti-PHB1 (Abcam, ab75766, 1:500), mouse anti-NDUFA13 (Santa Cruz Biotechnology, sc-66195, 1:1000), goat anti-TIM23 (Santa Cruz Biotechnology, sc-13298, 1:1000), rabbit anti-HSP60 (Cell Signaling Technology, #4870, 1:250), rabbit anti-Smac (Abcam, ab10919, 1:500), mouse anti-NDUFA9 (Abcam, ab14713, 1:500), mouse anti-NDUFS1 (Abcam, ab52690, 1:500), rabbit anti-OPA1 (Abcam, ab42364, 1:1000), mouse anti-VDAC (Abcam, ab14734, 1:1000), and rabbit anti-β-Actin (Cell Signaling Technology, #4970, 1:1000). Custom rabbit anti-acetyl-NDUFA13-K7 antibody was generated by immunizing rabbits with synthetic acetylated K7 peptide (ABclonal Technology, Wuhan, China).

### Cell Culture

Mouse embryonic fibroblasts (MEFs) were isolated from E13.5 embryos by standard procedures^39^. NDUFA13-deficient MEFs were generated by CRISPR/Cas9-mediated gene knockout. HEK293T cells, AC16 human cardiomyocytes, and H9C2 rat cardiomyoblasts were maintained in Dulbecco’s Modified Eagle Medium (DMEM; Corning) containing 4.5 g/L glucose, 4 mM L-glutamine, and 1 mM sodium pyruvate, supplemented with 10% fetal bovine serum (FBS; Life Technologies) and 1% penicillin-streptomycin at 37°C in a humidified atmosphere containing 5% CO₂. Neonatal mouse cardiomyocytes (NMCMs) were isolated from 1–3-day-old C57BL/6 mice, as previously described^40^, and cultured in the same medium. PHB2-deficient HEK293T cells were generated using the krCRISPR system as described below.

### Generation of NDUFA13-K7Q Conditional Knock-In Mice

Cardiomyocyte-specific NDUFA13-K7Q knock-in mice were generated using CRISPR/Cas9 technology (Cyagen Biosciences, Guangzhou, China). The mouse *Ndufa13* gene (NCBI Reference Sequence: NM_023312.3) is located on chromosome 8. A targeting cassette containing the loxP-flanked wild-type *Ndufa13* coding sequence with the K7Q point mutation (AAG→CAG) in exon 2 was inserted upstream of the endogenous start codon. Homology arms were then amplified from a BAC clone. Cas9 mRNA and guide RNA were co-injected with the targeting vector into C57BL/6N fertilised eggs. Founder mice were genotyped by PCR and verified by Sanger sequencing. Conditional knock-in mice (NDUFA13-K7Q^fl/fl^) were then crossed with Myh6-Cre transgenic mice to generate cardiomyocyte-specific NDUFA13-K7Q expression (Myh6^Cre+^; NDUFA13-K7Q^fl/fl^).

### Animal Procedures and Transverse Aortic Constriction

All animal procedures were approved by the Animal Ethics Review Committee of Zhejiang University (2023-1448) and performed in accordance with National Institutes of Health (NIH) guidelines. Mice were housed under controlled conditions (22 ± 1°C, 65–70% humidity, 12-hour light/dark cycle) with free access to food and water. Male mice aged 6–8 weeks underwent transverse aortic constriction (TAC) surgery to induce pressure overload-induced HF^41^. In brief, mice were anesthetized with isoflurane and a left thoracotomy was performed at the second intercostal space. A 7-0 silk suture was placed around the transverse aorta and tied against a 26-gauge needle to create stenosis. Sham-operated mice underwent identical procedures without aortic constriction. Mice were monitored daily and euthanised at the indicated time points for tissue collection. Animals were sacrificed at 4 weeks post-surgery for the K7Q knock-in characterisation experiments. For the validation cohort (Extended Data Fig. 3) and in vivo NR rescue experiment (Fig. 5h – l), tissues were collected at 6 weeks post-surgery.

### Human Cardiac Tissue Samples

Left ventricular tissue samples were obtained from patients with end-stage heart failure with reduced ejection fraction (HFrEF) undergoing heart transplantation. Control samples were obtained from donor hearts deemed unsuitable for transplantation. Tissues were immediately snap-frozen in liquid nitrogen and stored at -80°C. This study was conducted in accordance with the principles outlined in the Declaration of Helsinki. It received ethical approval from the Ethical Committee for Human Studies at The Second Affiliated Hospital of Zhejiang University School of Medicine (approval no. 2014-160). Clinical trial number: not applicable. All patients provided written informed consent.

### Echocardiography

Transthoracic echocardiography was performed using a Vevo 2100 system (FUJIFILM VisualSonics, Toronto, ON, Canada) equipped with a 30-MHz transducer. Mice were anesthetized with 1%–2% isoflurane and placed on a heated platform. M-mode images were obtained at the level of the papillary muscles to measure left ventricular internal diameter at end-diastole (LVIDd), left ventricular internal diameter at end-systole (LVIDs), and calculate fractional shortening (FS) and left ventricular mass. All measurements were averaged from at least three consecutive cardiac cycles.

### Mitochondrial Isolation

Mitochondria were isolated from cultured cells or cardiac tissues by standard procedures^42^. In brief, samples were homogenised in ice-cold isolation buffer (225 mM mannitol, 75 mM sucrose, 10 mM HEPES, 1 mM EGTA, 0.5% fatty acid-free BSA, pH 7.4) using a glass-Teflon homogeniser (10 strokes). Homogenates were centrifuged at 800 × g for 5 minutes at 4°C to remove nuclei and debris. The supernatant was then centrifuged at 8,000 × g for 10 minutes at 4°C to pellet mitochondria. The mitochondrial pellet was washed twice in BSA-free isolation buffer and resuspended at a concentration of 0.5–1.0 mg protein/mL. Protein concentration was then determined by the Bradford assay. Mitochondrial preparations were used within 2 hours of isolation.

### High-Resolution Respirometry

Mitochondrial oxygen consumption was measured using the Oxygraph-2k high-resolution respirometer (Oroboros Instruments, Innsbruck, Austria) at 37°C. Optimal concentrations of substrates and inhibitors were determined by preliminary titration experiments to ensure complete saturation or inhibition without non-specific effects. Isolated mitochondria (50–100 μg protein) were added to MiR05 respiration medium in each chamber. A substrate-uncoupler-inhibitor titration (SUIT) protocol was applied with sequential additions: glutamate/malate (10/2 mM) for complex I-linked respiration, ADP (2.5 mM) for OXPHOS capacity, succinate (10 mM) for complex I+II-linked respiration, rotenone (0.5 μM) to inhibit complex I, antimycin A (2.5 μM) to inhibit complex III, and ascorbate/TMPD (2/0.5 mM) with sodium azide (100 mM) for complex IV activity. Oxygen flux (pmol O₂ ·s⁻¹·mg⁻¹ protein) was recorded using DatLab software and normalised to mitochondrial protein content or mtDNA copy number when indicated.

### Submitochondrial Fractionation

For submitochondrial localisation analysis, isolated mitochondria were subjected to hypotonic swelling to selectively disrupt the outer mitochondrial membrane. Mitochondria were incubated in hypotonic buffer (20 mM HEPES, pH 7.4) for 30 minutes on ice, followed by treatment with or without proteinase K (50 μg/mL) for 30 minutes on ice. Proteinase K was inactivated by the addition of PMSF (2 mM). For complete solubilisation, mitochondria were treated with 1% Triton X-100. Samples were then analysed by western blotting with Smac (intermembrane space marker) and HSP60 (matrix marker) as compartment controls.

### Alkaline Carbonate Extraction

To distinguish integral membrane proteins from peripherally associated proteins, isolated mitochondria were subjected to alkaline extraction with 100 mM Na₂ CO₃ (pH 11.5) for 30 minutes on ice. Samples were centrifuged at 100,000 × g for 30 minutes at 4°C to separate membrane-integrated (pellet) and soluble (supernatant) fractions. TIM23 served as an integral inner mitochondrial membrane protein control; Smac served as a soluble protein control.

### Blue Native Polyacrylamide Gel Electrophoresis (BN-PAGE)

Isolated mitochondria were solubilised in BN-PAGE lysis buffer (20 mM Bis-Tris pH 7.0, 50 mM NaCl, 500 mM aminocaproic acid, 1 mM EDTA, 10% glycerol) containing 0.5% digitonin and protease inhibitors on ice for 15 minutes. Lysates were clarified by centrifugation at 20,000 × g for 20 minutes at 4°C. Supernatants were supplemented with 0.5% Coomassie Brilliant Blue G-250 and separated on 4%–16% gradient native gels (Invitrogen). Proteins were then transferred to PVDF membranes and immunoblotted with indicated antibodies.

### Immunoprecipitation and Western Blotting

Cells or tissues were lysed in IP lysis buffer (50 mM Tris-HCl pH 7.4, 150 mM NaCl, 1% NP-40, 0.5% sodium deoxycholate, protease inhibitors, and deacetylase inhibitors). For immunoprecipitation, lysates were incubated with primary antibodies overnight at 4°C with rotation, followed by incubation with Protein A/G Plus-Agarose beads (Santa Cruz Biotechnology) for 2 hours. Beads were washed five times with lysis buffer, and bound proteins were eluted by boiling in SDS sample buffer. For Western blotting, proteins were separated by SDS-PAGE, transferred to PVDF membranes, and probed with the indicated antibodies. Signals were detected using enhanced chemiluminescence (ECL) and quantified by ImageJ software (version 1.54f).

### GST Pulldown Assay

Recombinant GST-tagged PHB2 variants (wild-type, ΔPHB, ΔCC, K224Q/K236Q) and His-tagged NDUFA13 variants (wild-type, ΔTM, ΔCC, K7Q, K7R) were expressed in Escherichia coli BL21(DE3) and purified using glutathione-Sepharose 4B (GE Healthcare) or Ni-NTA agarose (Qiagen), respectively. For pulldown assays, GST-tagged proteins were immobilized on glutathione-Sepharose beads and incubated with His-tagged proteins in binding buffer (50 mM Tris-HCl pH 7.4, 150 mM NaCl, 0.1% Triton X-100) for 2 hours at 4℃. Beads were washed extensively, and bound proteins were analyzed by Western blotting.

### Mass Spectrometry Analysis

For identification of acetylation sites, immunoprecipitated NDUFA13 or PHB2 was separated by SDS-PAGE and excised gel bands were subjected to in-gel tryptic digestion. Peptides were analyzed by liquid chromatography-tandem mass spectrometry (LC-MS/MS) using an Orbitrap Fusion Lumos mass spectrometer (Thermo Fisher Scientific). Raw data were searched against the UniProt database using MaxQuant software (version 2.4.2.0) with acetylation (K) as a variable modification. For quantification of acetylation levels, parallel reaction monitoring (PRM) was performed using synthetic acetylated and non-acetylated peptide standards^43^.

### Interactome Analysis

For PHB2 interactome mapping, AC16 cells were treated with vehicle or oligomycin, lysed and subjected to immunoprecipitation using an anti-PHB2 antibody. Co-immunoprecipitated proteins were then identified by LC-MS/MS. Label-free quantification was performed using MaxQuant software (version 2.4.2.0). Differential interactors were then identified by volcano plot analysis with significance thresholds of fold change >2 and adjusted P < 0.05.

### Molecular Dynamics Simulations

The NDUFA13 structure was extracted from the cryo-EM structure of respiratory complex I (PDB: 6G2J)^18^. Missing N-terminal residues (1–4) were modelled using MODELLER (version 10.4)^44^. Acetylated K7 (AcK7) was parameterised using the Antechamber program (AmberTools 23) with RESP charges and GAFF force field^45,46^. Simulations were performed using GROMACS 2018.3 with the AMBER14SB force field for standard residues^47^. Proteins were solvated in a TIP3P water box with a minimum edge distance of 1.2 nm and neutralised with Na⁺ and Cl⁻ ions. Energy minimisation (5,000 steps, steepest descent) was followed by equilibration under NVT (100 ps) and NPT (100 ps) ensembles. Production simulations were run for 200 ns at 298 K and 1 bar using the V-rescale thermostat and Parrinello-Rahman barostat. A time step of 2 fs was used with LINCS constraints on all bonds. Long-range electrostatics were calculated using the particle mesh Ewald (PME) method. Trajectories were analysed for RMSD, radius of gyration (Rg), solvent-accessible surface area (SASA), hydrogen bonds, and secondary structure using GROMACS analysis tools and DSSP^48^.

### Free Energy Landscape Analysis

Free energy landscapes (FEL) were constructed from molecular dynamics trajectories using RMSD and radius of gyration as reaction coordinates. The free energy was calculated as ΔG = −k_B T ln(P), where P is the probability density of each bin. Representative structures were extracted from minimum energy basins for visualization.

### Calculation of Amino Acid Interaction Energy

During molecular dynamics simulations, protein conformations were recorded at regular intervals and the interaction energies between amino acid residues were calculated using the MM-GBSA (Molecular Mechanics-Generalised Born Surface Area) method. The total interaction energy between two amino acid residues within the protein monomer was expressed as: E_total = E_internal + E_elec + E_vdw + E_GB, where E_internal represented the bonded energy term defined by protein geometry, E_elec represented electrostatic interaction energy, E_vdw represented van der Waals interaction energy, and E_GB represented solvation energy accounting for the surrounding environment.

### Protein–Protein Docking and Binding Energy Calculations

Coiled-coil docking between NDUFA13 and PHB2 was performed using the HDOCK server with experimental restraints^49^. Multiple docking poses (parallel and antiparallel orientations) were evaluated. The protein complex system was constructed using CHARMM-GUI and solvated with TIP3P water molecules^50^. The simulation system was subjected to energy minimisation, followed by 200 ps NPT equilibration before calculating the interaction energy between protein components. Binding free energies were calculated using the MM/PBSA method implemented in the g_mmpbsa tool (version 1.0)^51^. Decomposition analysis was performed to identify key contributing residues. The error bars represent the standard deviation of the energy average obtained during the 200 ps NPT equilibration process.

### Transmission Electron Microscopy (TEM)

Cardiac tissues or cell pellets were fixed in 2.5% glutaraldehyde in 0.1 M phosphate buffer (pH 7.4) overnight at 4°C. Samples were post-fixed with 1% osmium tetroxide, stained with 2% uranyl acetate, dehydrated through a graded ethanol series, and embedded in epoxy resin. Ultrathin sections (70–80 nm) were cut using an ultramicrotome, mounted on copper grids, and counterstained with lead citrate. Images were subsequently acquired using a transmission electron microscope (Tecnai G2 Spirit, FEI). Mitochondrial area and cristae density were quantified using ImageJ software (version 1.54f). A minimum of 100 mitochondria per condition were analysed.

### Mitochondrial ROS Measurement

Mitochondrial reactive oxygen species (ROS) were measured using MitoSOX Red (Invitrogen). Cells were incubated with 5 μM MitoSOX Red in serum-free medium for 30 min at 37 °C, washed twice with PBS, and analysed by flow cytometry (BD FACSCanto II; excitation 510 nm, emission 580 nm). At least 10,000 events were acquired per sample, and data were analysed using FlowJo software (version 10.8.1).

### Immunofluorescence Microscopy

Cells were fixed with 4% paraformaldehyde for 15 minutes, permeabilised with 0.1% Triton X-100 for 10 minutes and then blocked with 5% BSA for 1 hour. Cells were incubated with primary antibodies overnight at 4°C, followed by fluorophore-conjugated secondary antibodies for 1 hour at room temperature. Nuclei were counterstained with DAPI. For mitochondrial co-localisation, cells were pre-stained with MitoTracker Deep Red prior to fixation. Images were acquired using a confocal microscope (Zeiss LSM 880).

### *In Vitro* Deacetylation Assay

To obtain endogenously hyperacetylated NDUFA13, neonatal mouse cardiomyocytes (NMCMs) were treated with oligomycin (OLG) to inhibit mitochondrial ATP synthase and induce OXPHOS dysfunction, as described above. Cells were harvested in IP lysis buffer supplemented with deacetylase inhibitors (10 mM nicotinamide and 1 μM trichostatin A) to preserve the acetylation state. Endogenous NDUFA13 was immunoprecipitated using anti-NDUFA13 antibody (Santa Cruz Biotechnology, sc-66195) cross-linked to Protein A/G magnetic beads (Thermo Fisher Scientific) at 4°C overnight, followed by three washes with IP wash buffer (50 mM Tris-HCl pH 7.4, 150 mM NaCl, 0.1% NP-40). The immunoprecipitated NDUFA13 was then eluted with 0.1 M glycine (pH 2.5) for 10 min at room temperature with gentle agitation, and the eluate was immediately neutralised by addition of 1 M Tris-HCl (pH 9.0) to restore physiological pH. The eluate was briefly centrifuged to remove residual beads, and the supernatant was used directly as the acetylated substrate in subsequent deacetylation reactions.

*In vitro* deacetylation reactions were performed in a total volume of 50 μL containing the eluted NDUFA13 substrate, 0.5 μg recombinant human SIRT3 protein (MedChemExpress, HY-P701609), and 500 μM β-NAD⁺ (Sigma-Aldrich) in deacetylation reaction buffer (50 mM Tris-HCl pH 8.0, 137 mM NaCl, 2.7 mM KCl, 1 mM MgCl_2_). Reactions were incubated at 37°C with gentle rotation and terminated at 0, 15, 30, 60, and 120 min by addition of 5× SDS loading buffer and immediate heating at 95°C for 5 min. Reaction products were resolved by SDS-PAGE and immunoblotted with the anti-acetyl-NDUFA13-K7 site-specific antibody, followed by re-probing with anti-NDUFA13 antibody as a loading control.

### krCRISPR System for Conditional Gene Knockout

To generate PHB2-deficient cells with rescue expression, we employed a dual episomal vector system (krCRISPR). The first plasmid expressed SpCas9 and tTA from the EF1α promoter via P2A self-cleaving peptides, with gRNA expressed from the U6 promoter. The second plasmid contained a TRE-driven cassette expressing copGFP, puromycin resistance, and the rescue gene (wild-type or mutant PHB2) with synonymous mutations in the gRNA target site. Co-transfection of both plasmids enabled simultaneous knockout of endogenous PHB2 and expression of exogenous variants.

### Histology

Hearts were fixed in 4% paraformaldehyde, embedded in paraffin, and sectioned at 5 μm thickness. Sections were stained with hematoxylin and eosin (H&E) for morphological assessment, picrosirius red for collagen deposition, and wheat germ agglutinin (WGA) conjugated to Alexa Fluor 488 for cardiomyocyte cross-sectional area measurement. Images were acquired using a light microscope or fluorescence microscope and analyzed using ImageJ software (version 1.54f).

### SIRT3 Deacetylase Activity Assay

SIRT3 deacetylase activity in cardiac tissues was assessed using a fluorometric SIRT3 Activity Assay Kit (Abcam, ab156067). Mitochondria were isolated from cardiac tissues of sham and TAC mice using a Tissue Mitochondria Isolation Kit, and protein concentration was determined by BCA assay. Twenty micrograms of mitochondrial protein per well was added to a 50 μL reaction volume according to the manufacturer’s instructions. Fluorescence was measured kinetically at 2-minute intervals over 30 minutes at room temperature (excitation 355 nm / emission 450 nm) using a fluorescence microplate reader. Each sample was assayed in technical duplicate and results were averaged. SIRT3 activity was expressed as relative fluorescence units normalized to protein input.

### *In Vivo* NR Rescue Experiment

To determine whether pharmacological restoration of NAD+ bioavailability could reverse NDUFA13-K7 acetylation-driven pathology in a K7-dependent manner, an *in vivo* rescue experiment was performed using four groups of mice: (i) WT + TAC + NS, (ii) WT + TAC + NR, (iii) K7Q knock-in + TAC + NS, and (iv) K7Q knock-in + TAC + NR (n = 7 per group). Beginning on day 1 post-TAC, mice in the NR groups received nicotinamide riboside (NR; 400 mg/kg/day) by intraperitoneal injection daily for 4 weeks; NS-treated control groups received an equal volume of 0.9% NaCl saline. Cardiac function was assessed by echocardiography at baseline and weekly for 6 weeks post-surgery, with blinded measurement of left ventricular fractional shortening (LVFS) and end-diastolic diameter (LVIDd). At 6 weeks post-surgery, animals were sacrificed and cardiac tissues were collected for molecular analyses. NDUFA13-K7 acetylation and PHB1/2 protein levels were assessed by immunoprecipitation and Western blotting as described above. PHB1/2 complex assembly was evaluated by blue native polyacrylamide gel electrophoresis (BN-PAGE). Mitochondrial ultrastructure and cristae density were quantified by transmission electron microscopy (TEM), with a minimum of 200 mitochondria per group analysed.

### Quantitative Real-Time PCR

Total DNA was extracted using a genomic DNA extraction kit (Qiagen). Mitochondrial DNA (mtDNA) content was quantified by real-time PCR using primers targeting the mitochondrial gene mt-Nd1 and normalized to the nuclear gene Hk2. Reactions were performed using PowerUp SYBR Green Master Mix (Applied Biosystems, Waltham, MA, USA) on a QuantStudio 6 Flex Real-Time PCR System (Applied Biosystems, Waltham, MA, USA).

### Cardiolipin Lipidomics by LC-MS/MS

Cardiac tissues (∼20 mg) were transferred to screw-cap tubes containing extraction solvent and homogenised using a tissue grinder (JXFSTPRP, Shanghai Jingxin) at 4°C and 64 Hz for three cycles of 5 minutes each. Homogenates were transferred to glass tubes, and lipids were extracted by addition of two volumes of chloroform, followed by vortexing for 1 minute and centrifugation at 1,000 × g for 5 minutes at room temperature. The supernatant was transferred to a new glass tube and centrifuged again under identical conditions. A 200 µL aliquot of the final supernatant was evaporated to dryness under vacuum at room temperature and stored at −80°C until analysis.

Prior to injection, dried lipid extracts were reconstituted in 200 µL of LC-MS-grade isopropanol:acetonitrile:water (2:1:1, v/v/v), vortexed for 10 minutes, and allowed to stand at room temperature for 30 minutes. Samples were centrifuged at 15,000 × g for 15 minutes at room temperature, and the supernatant was transferred to a new tube and centrifuged again at 15,000 × g for 15 minutes at 4°C. The final supernatant was transferred to MS vials for analysis.

Untargeted cardiolipin profiling was performed on a Thermo Scientific Orbitrap Exploris 240 (OE240) mass spectrometer coupled to a Waters ACQUITY UPLC system. Chromatographic separation was achieved on an ACQUITY UPLC BEH C18 column (1.7 µm, 2.1 × 50 mm) with a BEH C18 Vanguard pre-column at 45°C. The injection volume was 2 µL and the flow rate was 0.3 mL/min. Mobile phase A consisted of methanol:acetonitrile:water (1:1:1, v/v/v) containing 10 mM ammonium acetate and 0.1% formic acid; mobile phase B consisted of isopropanol:acetonitrile (9:1, v/v) containing 10 mM ammonium acetate. Gradient elution was performed as follows: 0–1.5 min, 32% B; 1.5–4 min, 32–45% B; 4–5 min, 45–52% B; 5–8 min, 52–58% B; 8–11 min, 58–66% B; 11–14 min, 66–70% B; 14–18 min, 70–75% B; 18–21 min, 75–97% B; 21–25 min, 97% B; 25.01 min, return to 32% B.

Mass spectrometric data were acquired in negative ion mode using heated electrospray ionisation (H-ESI) with a spray voltage of −2,500 V, sheath gas flow of 40 arb, and ion transfer tube temperature of 320°C. Full-scan spectra were acquired over m/z 500–1,800 at an Orbitrap resolution of 120,000 with RF lens at 70%. Data-dependent MS² (ddMS²) fragmentation was performed at an Orbitrap resolution of 15,000 using higher-energy collisional dissociation (HCD) at normalised collision energies of 30%, 40%, and 50%.

Raw data were processed using MS-DIAL software^52^. Cardiolipin species were annotated by matching precursor masses and MS² fragmentation spectra against both public lipid databases and an in-house cardiolipin spectral library, with [M−H]⁻ and [M−2H]²⁻ adducts considered. Peak intensities were normalised to the total cardiolipin signal intensity per sample to account for differences in sample loading.

### Targeted Metabolomics

Standard compounds were purchased from Sigma-Aldrich, Steraloids Inc., and TRC Chemicals. Stock solutions (5.0 mg/mL) were prepared in appropriate solvents and mixed to generate calibration standards. Cardiac tissue samples (∼10 mg) were thawed on ice, homogenised with 20 μL deionised water and zirconia beads for 3 min, followed by addition of 120 μL pre-cooled methanol containing internal standards and homogenisation for another 3 min. Samples were centrifuged at 18,000 g for 20 min at 4°C, and 20 μL supernatant was transferred to a 96-well plate. Derivatisation was performed using an Eppendorf epMotion workstation: 20 μL freshly prepared derivatisation reagent was added to each well, incubated at 30°C for 60 min, followed by addition of 330 μL pre-cooled 50% methanol. After centrifugation (4,000 g, 30 min, 4°C), 135 μL supernatant was transferred to a new plate with 10 μL internal standard.

Metabolites were quantified by ultra-high-performance liquid chromatography-tandem mass spectrometry (UPLC-MS/MS; ACQUITY UPLC-Xevo TQ-S, Waters Corp.). Raw data were processed using QuanMET software (version 2.0, Metabo-Profile) for peak integration, calibration, and quantification.

### Statistical Analysis of Metabolomics Data

Differential metabolites were identified using orthogonal partial least squares discriminant analysis (OPLS-DA) with variable importance in projection (VIP) > 1.0 as the threshold. Univariate analysis (t-test or Mann-Whitney U test, selected based on data distribution) was performed in parallel. Potential biomarkers were identified by intersecting results from multivariate and univariate analyses. Pathway enrichment analysis was performed using the Small Molecule Pathway Database (SMPDB)^53^ and Predicted Metabolite Sets libraries via MetaboAnalyst 5.0^54^. Statistical analyses were conducted using R (version 4.2.0).

### Statistical Analysis

Data are presented as mean ± SD as indicated in figure legends. Statistical analyses were performed using GraphPad Prism (version 9.5.1). Normality was assessed using the Kolmogorov-Smirnov test. Comparisons between two groups were performed using two-tailed unpaired Student’s t-tests. Comparisons between multiple groups were performed using one-way or two-way analysis of variance (ANOVA) followed by Bonferroni’s post hoc test. Survival curves were analysed using the Kaplan-Meier method and compared by the log-rank test. P < 0.05 was considered statistically significant. *P < 0.05; **P < 0.01; ***P < 0.001; ****P < 0.0001; ns, not significant.

---

**Extended Fig. 1.**
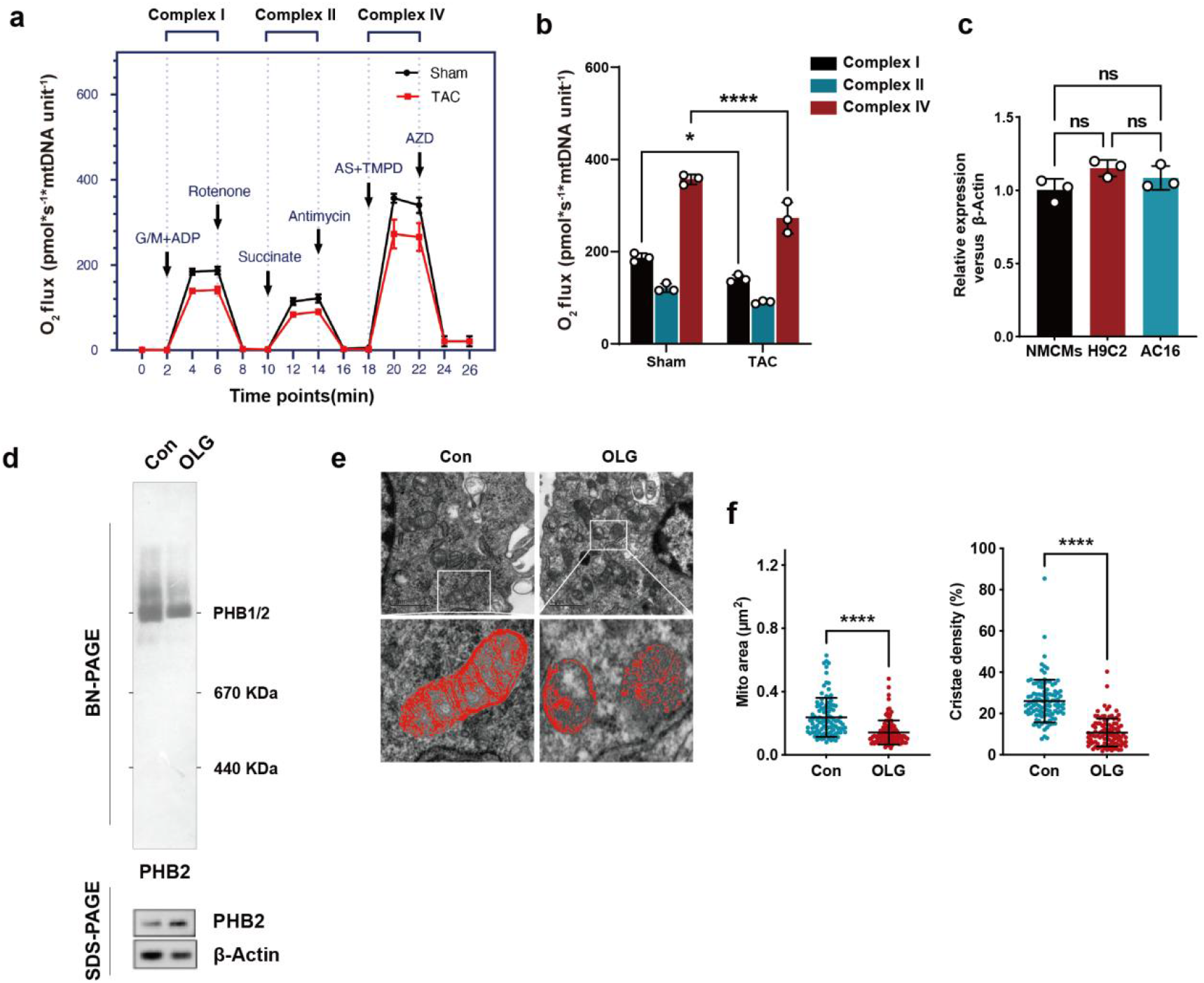
|. OXPHOS Inhibition Induces PHB1/2 Disassembly and Cristae Loss. **a,** Mitochondrial oxygen consumption rates in isolated cardiac mitochondria from sham-operated (Sham) and transverse aortic constriction (TAC) mice, as measured by high-resolution respirometry (Oroboros O2k). Oxygen flux was normalized to mitochondrial DNA (mtDNA) content. A substrate-uncoupler-inhibitor titration protocol was applied with sequential additions of substrates (glutamate/malate + ADP, succinate, ascorbate/TMPD) and inhibitors (rotenone, antimycin A, sodium azide) to assess complex I, II, and IV activities. **b,** Quantification of oxygen flux. n = 3 mice per group. **c,** Relative mitochondrial DNA (mtDNA) content assessed by quantitative PCR (qPCR) in neonatal mouse cardiomyocytes (NMCMs), H9C2 cells, and AC16 cells following indicated treatments, normalized to nuclear DNA content. **d,** BN-PAGE analysis of PHB1/2 complex assembly in AC16 cells treated with vehicle or oligomycin (OLG). **e-f,** Representative TEM images (**e**) of mitochondrial ultrastructure in AC16 cells treated with vehicle or OLG, with quantification of cristae density (**f**). Scale bar, 1 μm. n = 100 mitochondria per group (5 images × 5 mitochondria per image, 4 mice per group). Data are presented as mean ± SD from one representative experiment of at least three independent experiments. Statistical significance was determined by two-way analysis of variance (ANOVA) with Bonferroni’s post hoc test (**b**), one-way ANOVA (**c**), and two-tailed unpaired Student’s t-test (**f**). ***P < 0.001; ****P < 0.0001; ns, no significance.

**Extended Fig. 2.**
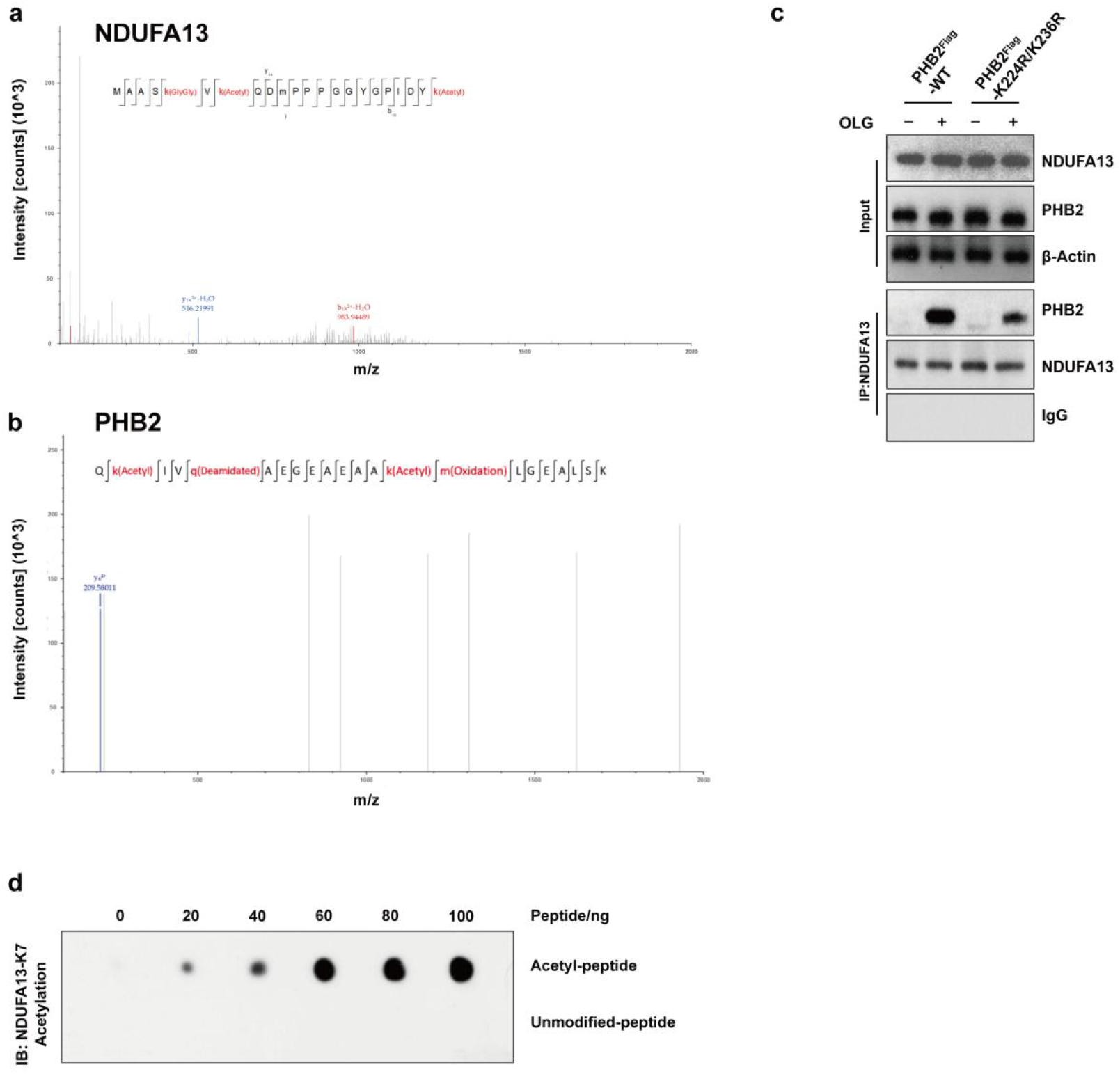
|. Identification and Validation of Acetylation Sites on NDUFA13 and PHB2. **a,** Identification of NDUFA13 acetylation sites by immunoprecipitation followed by mass spectrometry analysis. Representative mass spectrum showing the K7 acetylation site on NDUFA13. **b,** Identification of PHB2 acetylation sites by immunoprecipitation followed by mass spectrometry analysis. Representative mass spectrum showing acetylation modifications at K224 and K236 on PHB2. **c,** Co-immunoprecipitation (Co-IP) and Western blot analysis of NDUFA13-PHB2 interaction in PHB2-deficient HEK293T cells reconstituted with Flag-tagged PHB2-WT or PHB2-K224R/K236R following OLG treatment. **d,** Validation of a custom anti-acetyl-NDUFA13-K7 antibody specificity using dot blot assay with acetylated and non-acetylated K7 peptides. Blots shown are representative of at least three independent experiments.

**Extended Fig. 3.**
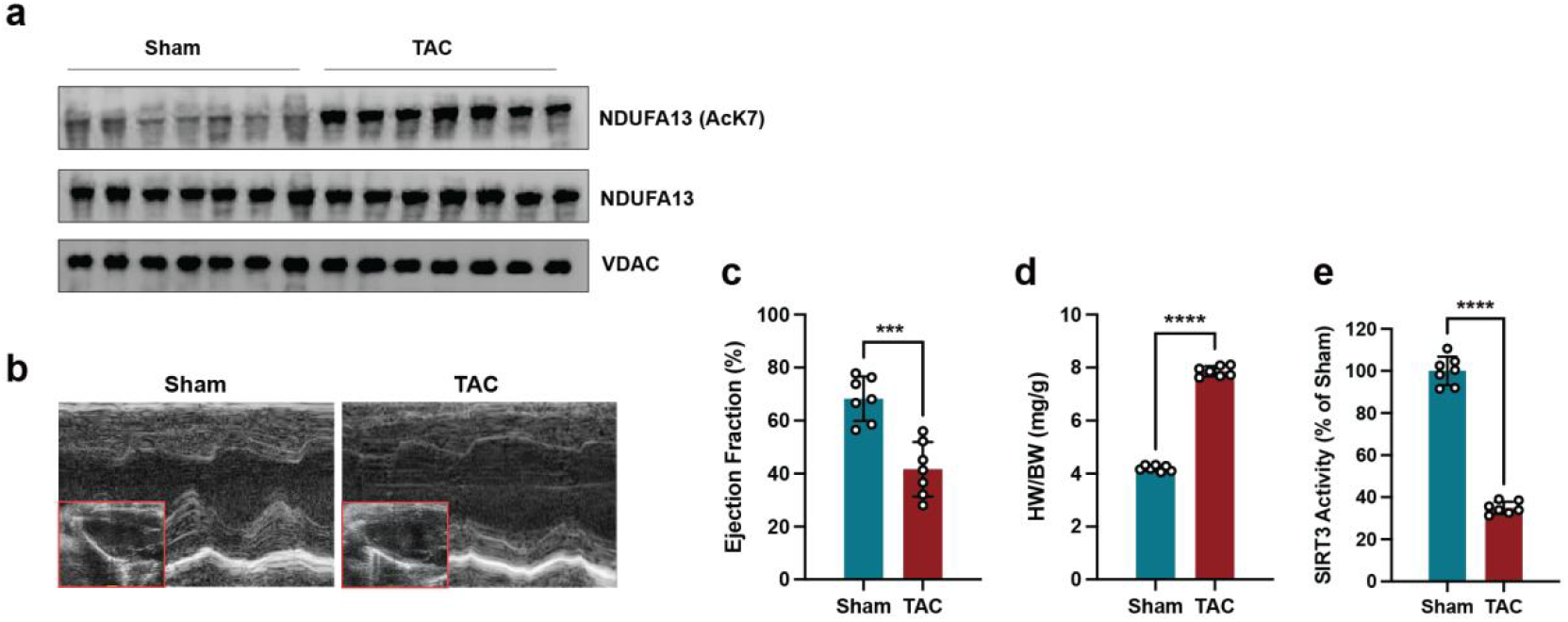
|. Validation of NDUFA13-K7 Hyperacetylation and SIRT3 Inactivation in a Mouse Model of Pressure Overload-Induced Heart Failure. **a**, NDUFA13-K7 acetylation in pressure overload-induced heart failure. IP/WB showing AcK7 (upper), total NDUFA13 (middle), and VDAC loading control (lower) in sham and TAC cardiac lysates. **b,** Representative M-mode echocardiographic images of the left ventricle from indicated experimental groups. **c-d,** Quantification of ejection fraction (**c**) and heart weight/body weight ratio (**d**) is shown. n = 7 mice per group. **e,** SIRT3 deacetylase activity in cardiac tissues from Sham and TAC mice, assessed by fluorometric assay. Data are presented as mean ± SD from one representative experiment of at least three independent experiments. Statistical significance was determined by two-tailed unpaired Student’s t-test (**c-e**). ***P < 0.001; ****P < 0.0001.

**Extended Fig. 4.**
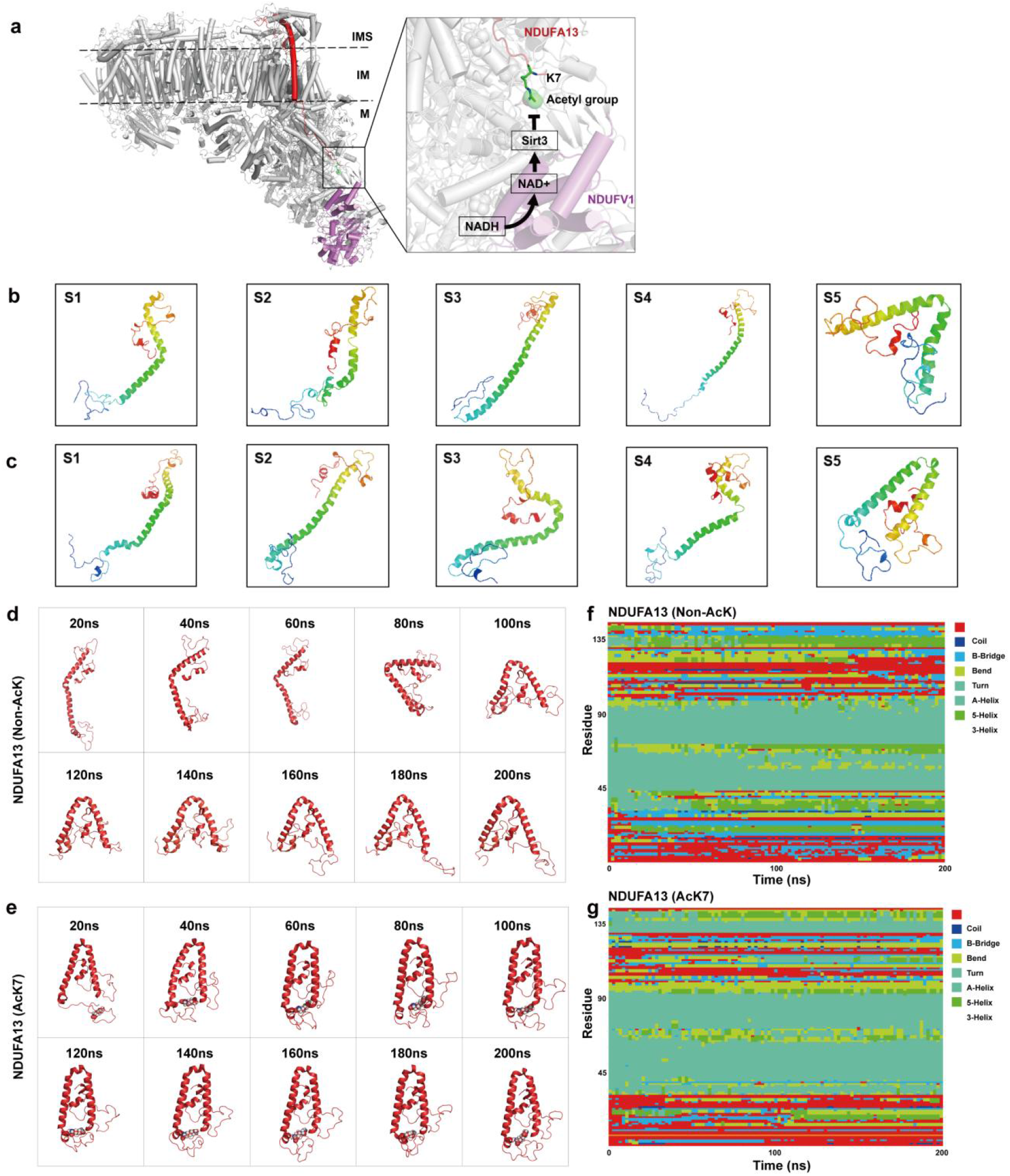
|. Structural Positioning of NDUFA13-K7 and Conformational Dynamics Upon Acetylation. **a,** Structural visualization of NDUFA13-K7 positioning within the NADH dehydrogenase complex (respiratory chain complex I) based on the cryo-EM structure (PDB: 6G2J). The K7 residue is located in proximity to the NADH binding site, suggesting its capacity to sense OXPHOS functional status through acetylation. **b-c,** Free energy landscape (FEL) analysis showing global energy minima (ΔG, kJ/mol) of acetylated NDUFA13 (AcK7) and non-acetylated NDUFA13 (Non-AcK7) as a function of RMSD (nm) and radius of gyration (Rg, nm) over 200 ns molecular dynamics simulations. Representative conformations extracted from the minimum energy basins are shown for AcK7 (**b**) and Non-AcK7 (**c**). **d-e,** Snapshots of NDUFA13 conformations extracted at indicated time points during 200 ns molecular dynamics simulations, illustrating the dynamic structural evolution of non-acetylated (Non-AcK7, **d**) and acetylated (AcK7, **e**) NDUFA13. **f-g**, Secondary structure analysis of non-acetylated (Non-AcK7, **f**) and acetylated (AcK7, **g**) NDUFA13 over the 200 ns simulation trajectory. Secondary structure elements including coil, β-bridge, bend, turn, α-helix, 3₁ ₀-helix (3-helix), and π-helix (5-helix) are color-coded as indicated in the legend.

**Extended Fig. 5.**
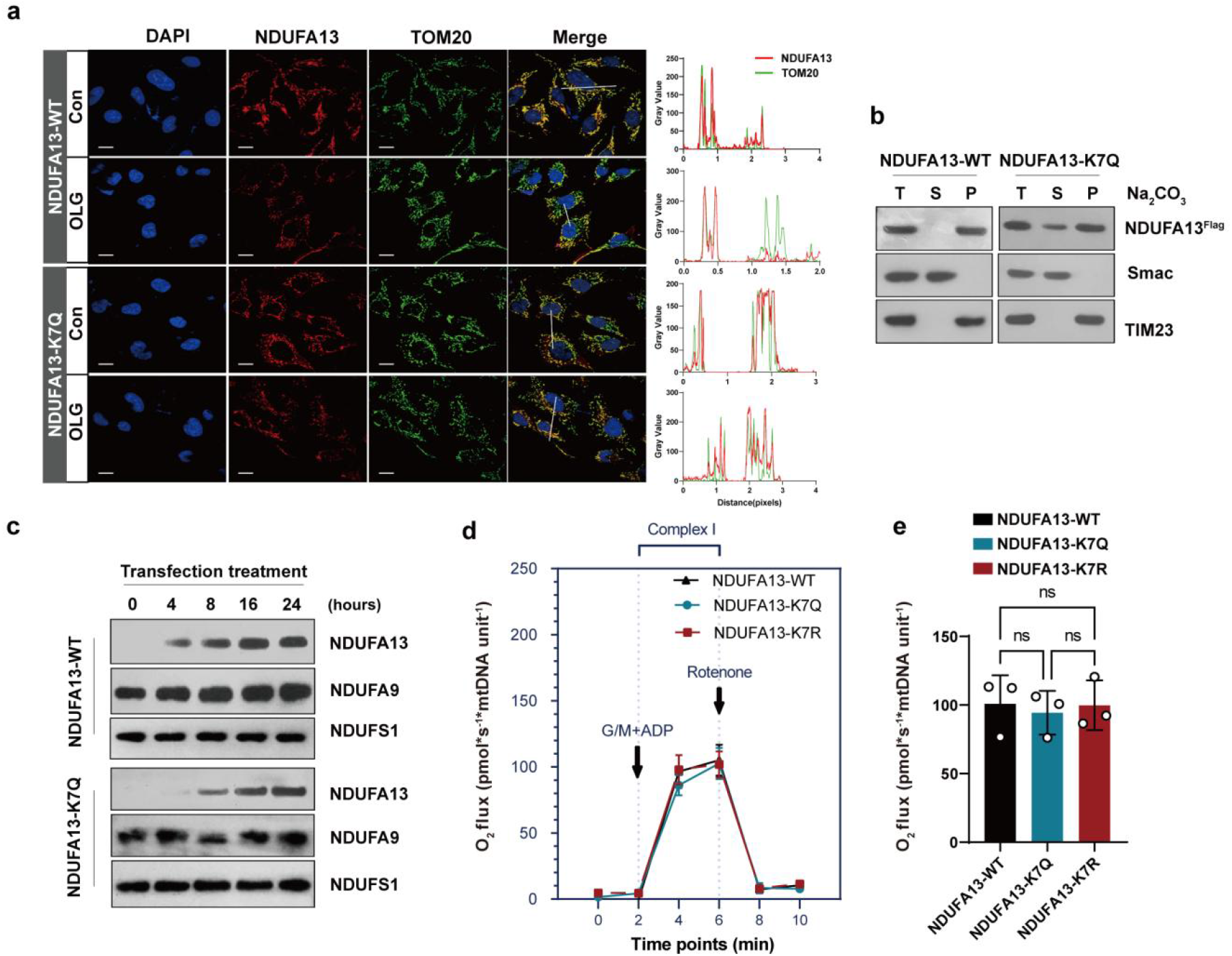
|. Subcellular Localization, Membrane Integration, and Functional Characterization of NDUFA13 Variants. **a,** Immunofluorescence microscopy analysis of NDUFA13 subcellular localization in NDUFA13-deficient MEF cells reconstituted with indicated NDUFA13 variants (WT, K7Q, or K7R) under vehicle or OLG-treated conditions. Scale bar, 10 μm. **b,** Membrane integration analysis of NDUFA13 variants in NDUFA13-deficient MEF cells. Isolated mitochondria were subjected to alkaline extraction with Na₂ CO₃ (pH 11.5), followed by centrifugation to separate membrane-integrated (P, pellet) and soluble (S, supernatant) fractions. TIM23 (integral inner mitochondrial membrane protein) and Smac (soluble intermembrane space protein) served as fractionation controls. T, total lysate. **c,** Quantitative analysis of NDUFA13-WT and NDUFA13-K7Q incorporation rates into complex I at indicated time points following transfection into NDUFA13-deficient MEF cells. **d-e,** Mitochondrial oxygen consumption rates in isolated mitochondria from NDUFA13-deficient MEF cells reconstituted with wild-type (WT), K7Q, or K7R NDUFA13, measured by high-resolution respirometry (Oroboros O2k). Oxygen flux was normalized to mitochondrial DNA (mtDNA) content. Complex I activity was assessed using glutamate/malate as substrates. n = 3 independent experiments (**d**). Quantification of oxygen flux was shown (**e**). Data are presented as mean ± SD from one representative experiment of at least three independent experiments. Statistical significance was determined by one-way ANOVA (**e**) with multiple comparison. ns, no significance.

**Extended Fig. 6.**
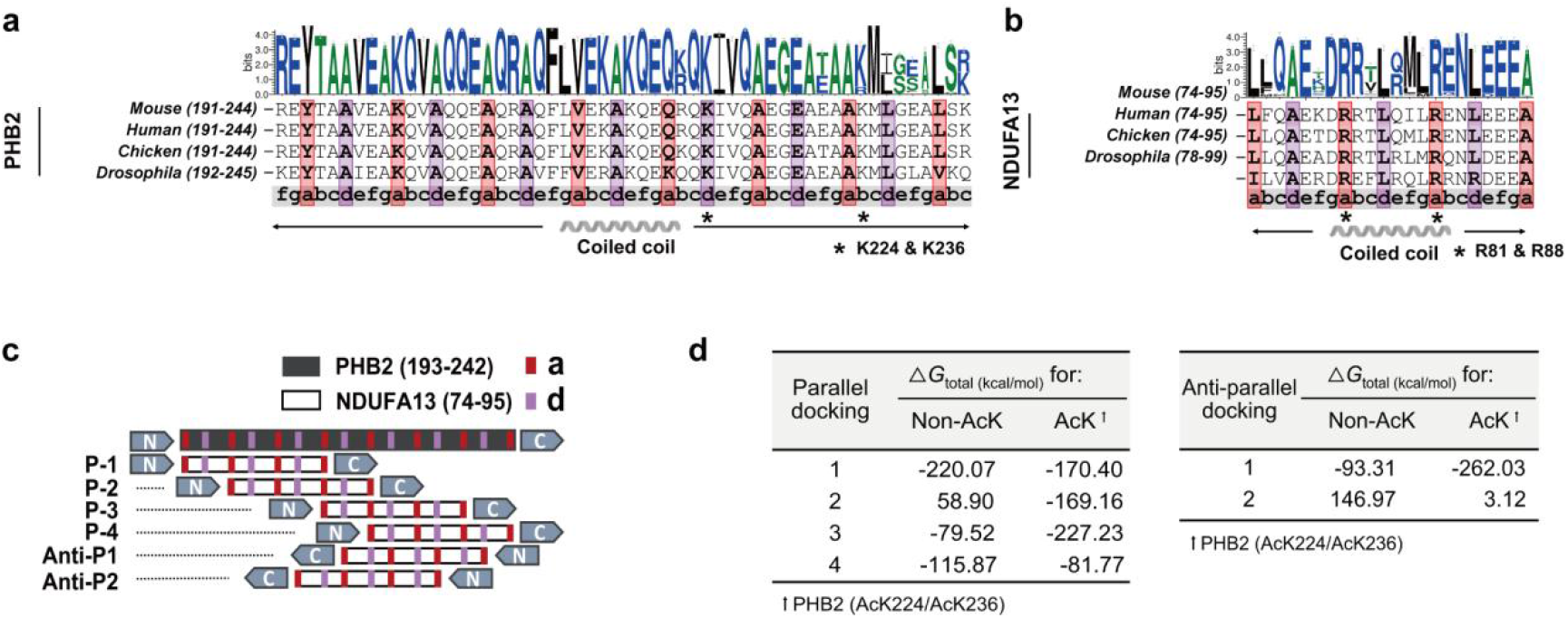
|. Sequence Conservation and Docking Analysis of NDUFA13-PHB2 Coiled-Coil Interactions. **a-b,** Sequence logo and multiple sequence alignment of heptad repeat residues within the coiled-coil (CC) domains of PHB2 (**a**) and NDUFA13 (**b**) across species. Residues at the “a” and “d” positions critical for knobs-into-holes (KIH) packing are highlighted. **c,** Schematic model illustrating potential coiled-coil interaction modes between NDUFA13 and PHB2 based on KIH packing principles. Four parallel and two antiparallel docking orientations are shown. **d,** Free binding energy analysis of NDUFA13-PHB2 coiled-coil interactions. Tables display calculated docking energies for parallel (left) and antiparallel (right) orientations, comparing non-acetylated and acetylated (K224/K236) PHB2.

**Extended Fig. 7.**
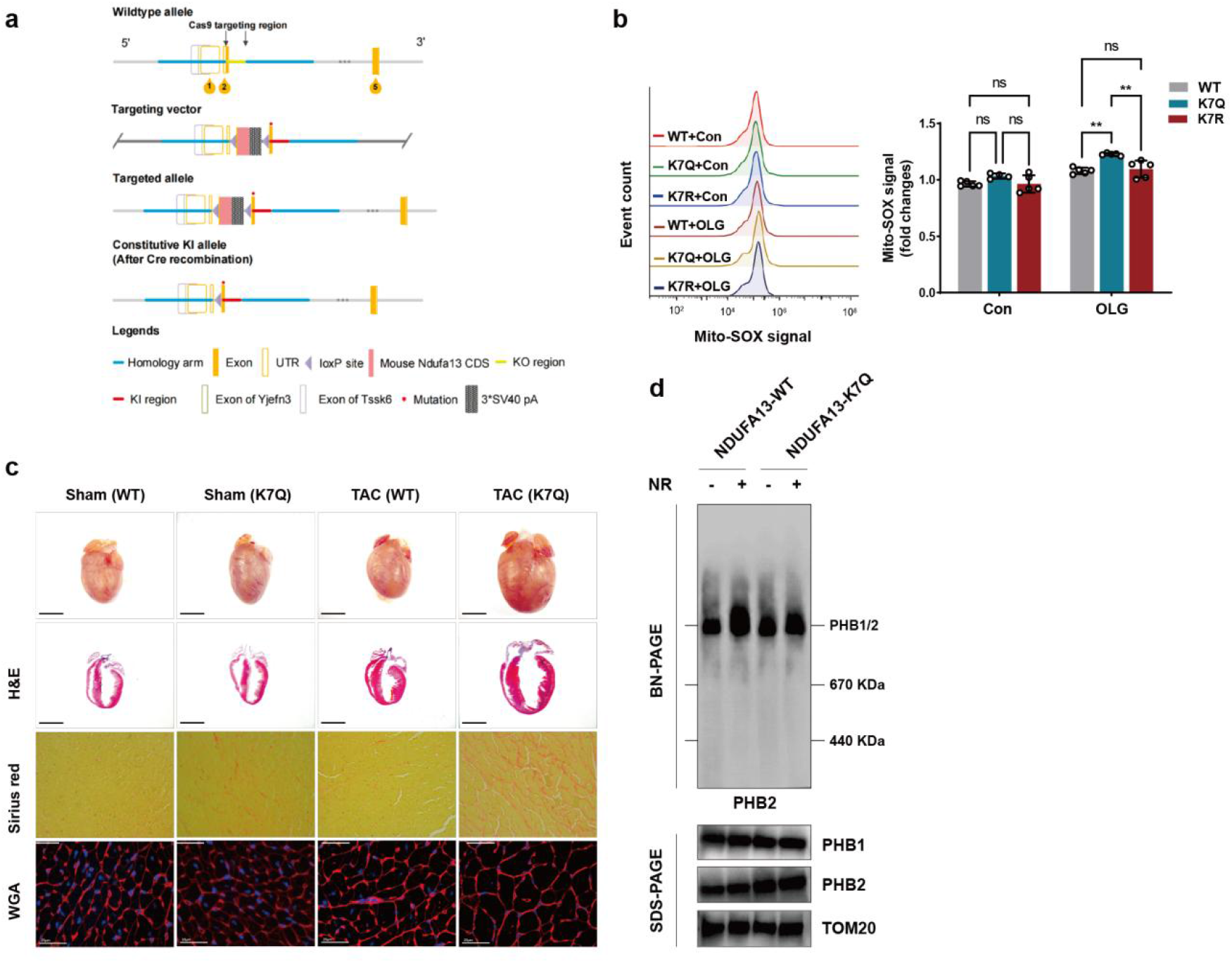
|. Characterization of the NDUFA13-K7Q Knock-In Mouse Model and Molecular Validation of *In Vivo* NR Rescue. **a,** Schematic of the CRISPR/Cas9-mediated knock-in strategy for generating NDUFA13-K7Q conditional knock-in mice. The mouse Ndufa13 gene (NCBI Reference Sequence: NM_023312.3) is located on chromosome 8 and contains five exons, with the ATG start codon in exon 2 and the TAG stop codon in exon 5. The targeting cassette (loxP–5′UTR–mouse Ndufa13 CDS–3×SV40 pA–loxP) was inserted 7–10 bp upstream of the ATG start codon. The K7Q point mutation (AAG→CAG) was introduced into exon 2 within the knock-in region. Homology arms were amplified by PCR using a BAC clone as template. Cas9 mRNA and gRNA were co-injected with the targeting vector into fertilized eggs. Founder pups were genotyped by PCR and verified by Sanger sequencing. **b,** Flow cytometric analysis of mitochondrial reactive oxygen species (ROS) production using MitoSOX fluorescence in NDUFA13-deficient MEF cells expressing the indicated NDUFA13 variants following OLG treatment (left), with corresponding quantification (right). **c,** Representative histological images of cardiac tissues from indicated experimental groups. Sections were stained with hematoxylin and eosin (H&E; scale bar, 3 mm), picrosirius red for fibrosis assessment (scale bar, 100 μm), and wheat germ agglutinin (WGA) for cardiomyocyte cross-sectional area determination (scale bar, 25 μm). **d,** BN-PAGE analysis of PHB1/2 assembly in cardiac tissues from WT and NDUFA13-K7Q knock-in mice treated with NR or vehicle following TAC. Data are presented as mean ± SD from one representative experiment of at least three independent experiments. Statistical significance was determined by two-way analysis of variance (ANOVA) with Bonferroni’s post hoc test (**b**). **P < 0.01; ns, no significance.

**Extended Fig. 8.**
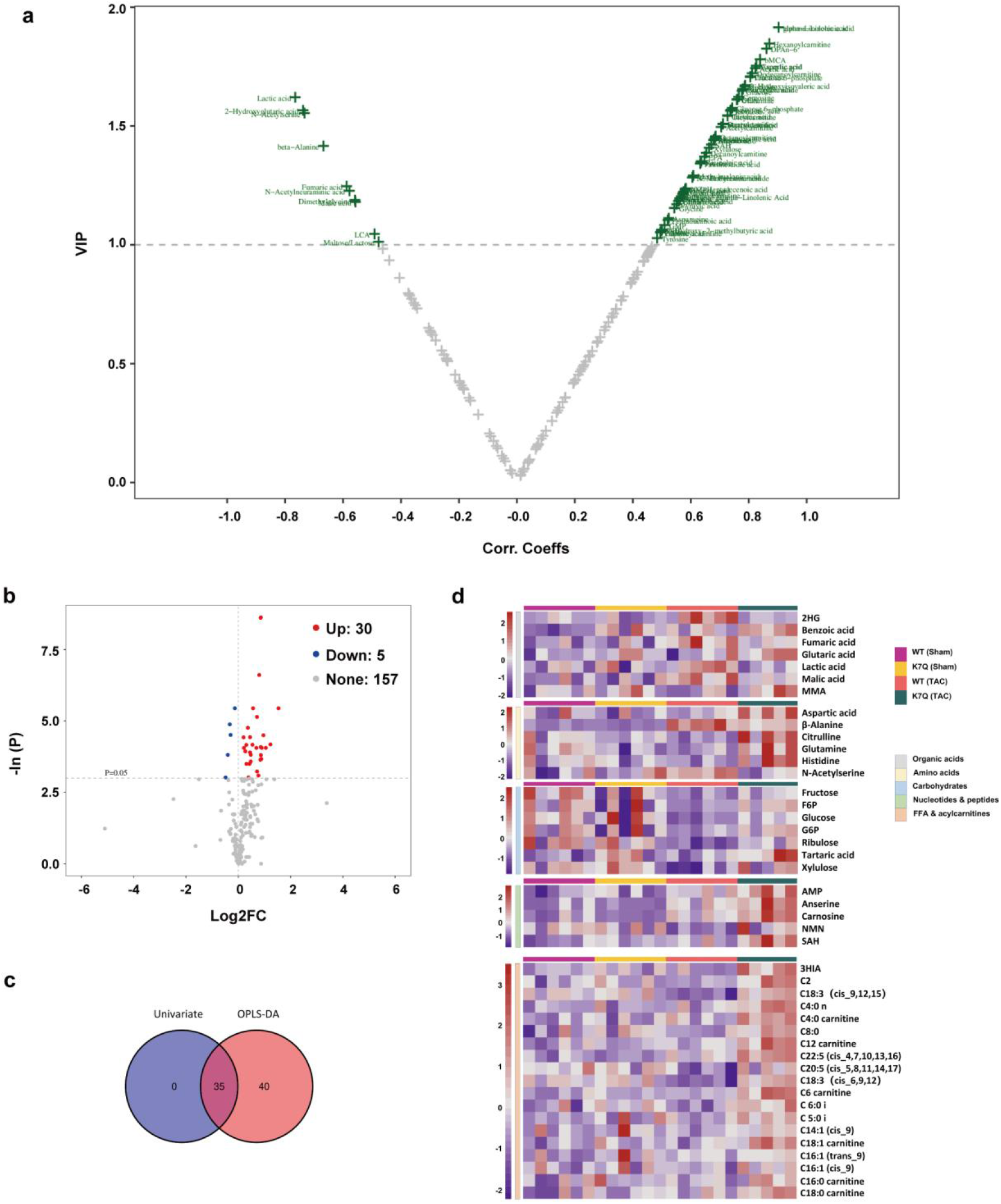
|. Metabolomic profiling and differential metabolite analysis in NDUFA13-K7Q knock-in hearts. **a,** Volcano plot of differential metabolites identified by orthogonal partial least squares discriminant analysis (OPLS-DA). **b,** Volcano plot of differentially abundant metabolites based on univariate analysis. Red points indicate upregulated metabolites; blue points indicate downregulated metabolites. **c,** Venn diagram showing overlap of differential metabolites identified by univariate and multivariate (OPLS-DA) statistics. **d,** Heatmap showing differential metabolite profiles between cardiomyocyte-specific NDUFA13-K7Q knock-in mice (Myh6^Cre+^; NDUFA13-K7Q^fl/fl^) and control littermates (Myh6^Cre−^; NDUFA13-K7Q^fl/fl^) following sham or TAC surgery.

**Extended Fig. 9.**
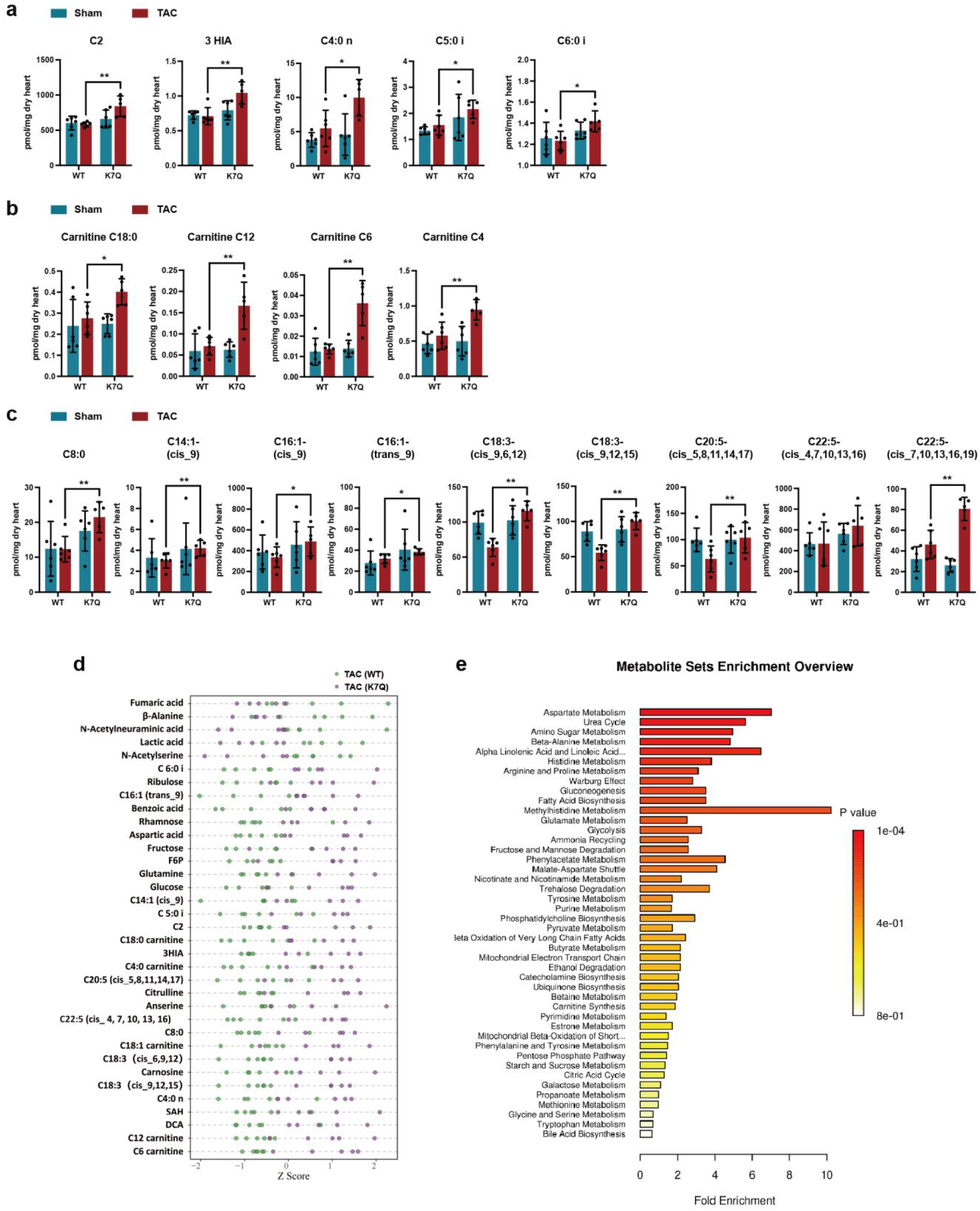
|. Impaired fatty acid oxidation and pathway alterations in NDUFA13-K7Q hearts. **a-c,** Absolute myocardial levels of dysregulated FFAs and acylcarnitine metabolites in the studied groups. **d,** Differential metabolite Z-score point plot demonstrating metabolic variation between NDUFA13-WT and NDUFA13-K7Q groups post-TAC surgery. The plot’s vertical axis lists the detected metabolites, while the horizontal axis shows the Z-score of differential metabolites. **e,** The bar chart represents the results of pathway enrichment analysis between NDUFA13-K7Q and NDUFA13-WT in the pathway-associated metabolite sets (SMPDB) database. The length of each bar on the x-axis corresponds to the fold enrichment, and the color of the bars represents the associated P-value. The intensity of the red color is positively correlated with the significance of the pathway. Data are presented as mean ± SD. Statistical significance was determined by two-tailed unpaired Student’s t-test (**a-c**). *P < 0.05; **P < 0.01.

**Extended Fig. 10.**
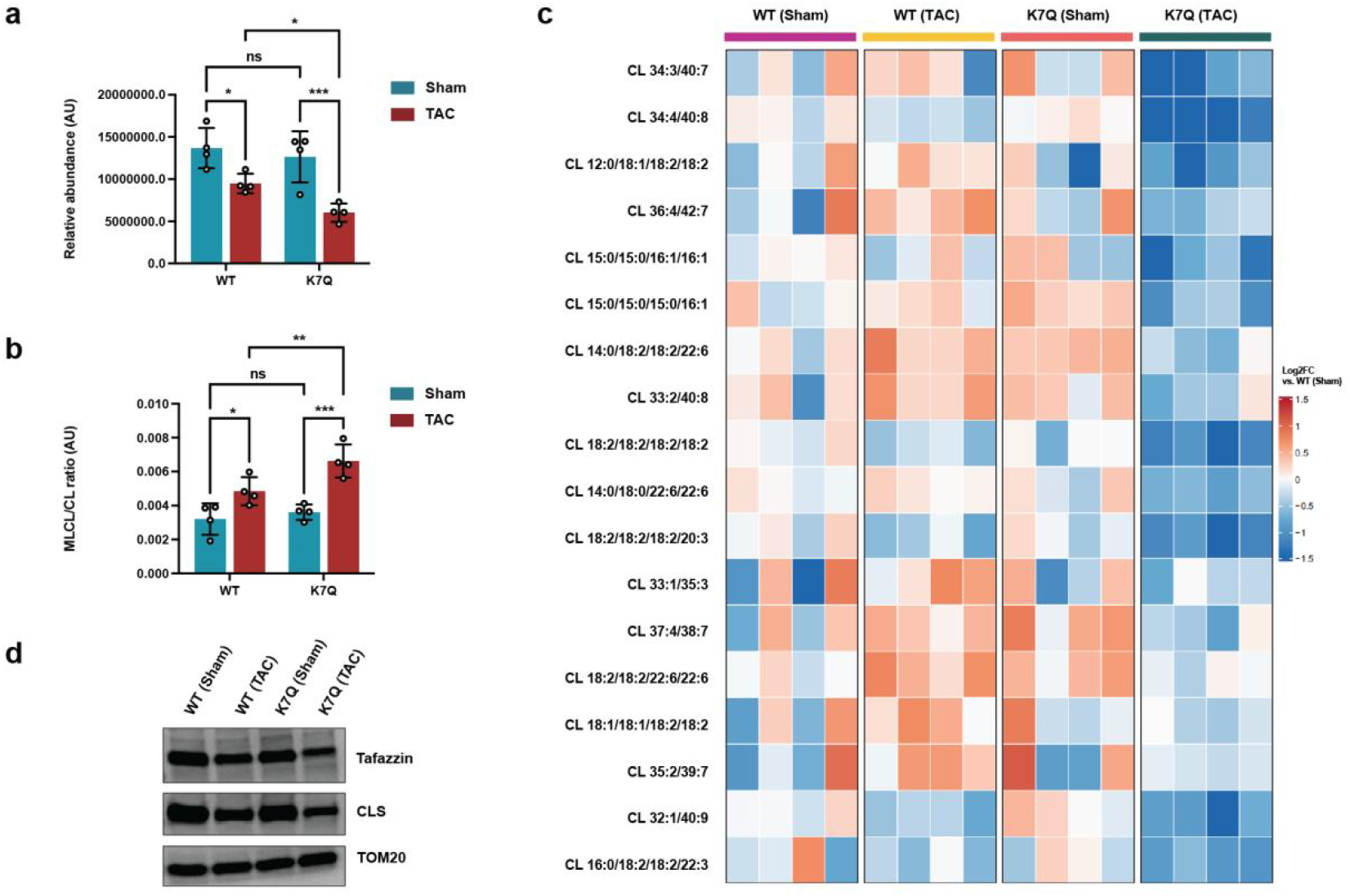
|. Cardiolipin Lipidomics Profiling Reveals Stress-Dependent CL Depletion in NDUFA13-K7Q Knock-In Hearts. **a,** Bar graph showing relative abundance of tetralinoleoyl-cardiolipin (L4CL; CL 18:2/18:2/18:2/18:2), the predominant cardiac cardiolipin species, in cardiac tissues from WT (Sham), WT (TAC), K7Q (Sham), and K7Q (TAC) mice. **b,** Bar graph showing the MLCL/CL ratio across the four experimental groups. MLCL (monolysocardiolipin) is a deacylation intermediate that accumulates when tafazzin-mediated cardiolipin reacylation is impaired. **c,** Heatmap showing the 18 cardiolipin species significantly decreased in K7Q (TAC) hearts relative to WT+TAC (all p < 0.05, one-way ANOVA with Bonferroni’s post hoc test). Each column represents an individual biological replicate (n = 4 per group). Colour scale reflects log2Fold change relative to the WT (Sham) group mean. Species are ordered by fold change in K7Q+TAC versus WT+TAC, with the most depleted species at the top. **d,** Representative Western blot showing protein levels of tafazzin (TAZ) and cardiolipin synthase (CLS) in cardiac tissues from the four experimental groups. TOM20 serves as a mitochondrial loading control. Data are presented as mean ± SD. Statistical significance was determined by two-way ANOVA with multiple comparison (a, b). *P < 0.05; **P < 0.01; ***P < 0.001; ns, no significance.

## References

1 Savarese, G. et al. Global burden of heart failure: a comprehensive and updated review of epidemiology. Cardiovasc. Res. 118, 3272–3287 (2023).

2 Feng, J., Zhang, Y. & Zhang, J. Epidemiology and Burden of Heart Failure in Asia. *JACC*. Asia 4, 249–264 (2024).

3 Teng, T. K. & Lam, C. S. P. Comprehensive Introduction to ASIAN-HF Registry. *JACC*. Asia 5, 1206–1216 (2025).

4 Brown, D. A. et al. Expert consensus document: Mitochondrial function as a therapeutic target in heart failure. Nat. Rev. Cardiol. 14, 238–250 (2017).

5 Lopaschuk, G. D., Karwi, Q. G., Tian, R., Wende, A. R. & Abel, E. D. Cardiac Energy Metabolism in Heart Failure. Circ. Res. 128, 1487–1513 (2021).

6 Karamanlidis, G. et al. Mitochondrial complex I deficiency increases protein acetylation and accelerates heart failure. Cell Metab. 18, 239–250 (2013).

7 Zhou, B. & Tian, R. Mitochondrial dysfunction in pathophysiology of heart failure. J. Clin. Invest. 128, 3716–3726 (2018).

8 Bock, F. J. & Tait, S. W. G. Mitochondria as multifaceted regulators of cell death. Nat. Rev. Mol. Cell Biol. 21, 85–100 (2020).

9 Merkwirth, C. & Langer, T. Prohibitin function within mitochondria: essential roles for cell proliferation and cristae morphogenesis. Biochim. Biophys. Acta 1793, 27–32 (2009).

10 Wang, C. et al. Krüppel like factor 7 regulates mitochondrial dynamics balance in myocardial infarction. *Commun*. Biol. 8, 722 (2025).

11 Rose, K. et al. In situ cryo-ET visualization of mitochondrial depolarization and mitophagic engulfment. Proc. Natl. Acad. Sci. U. S. A. 122, e2511890122 (2025).

12 Chen, L., Gong, Q., Stice, J. P. & Knowlton, A. A. Mitochondrial OPA1, apoptosis, and heart failure. Cardiovasc. Res. 84, 91–99 (2009).

13 Wai, T. et al. Imbalanced OPA1 processing and mitochondrial fragmentation cause heart failure in mice. Science 350, aad0116 (2015).

14 Lee, C. F. et al. Normalization of NAD+ Redox Balance as a Therapy for Heart Failure. Circulation 134, 883–894 (2016).

15 Horton, J. L., et al. Mitochondrial protein hyperacetylation in the failing heart. JCI insight 2 (2016).

16 Fiedorczuk, K. & Sazanov, L. A. Mammalian Mitochondrial Complex I Structure and Disease-Causing Mutations. Trends Cell Biol. 28, 835–867 (2018).

17 Shao, M. et al. Activation of RXRα exerts cardioprotection through transcriptional upregulation of Ndufs4 in heart failure. Sci. Bull. 69, 1202–1207 (2024).

18 Agip, A. A. et al. Cryo-EM structures of complex I from mouse heart mitochondria in two biochemically defined states. Nat. Struct. Mol. Biol. 25, 548–556 (2018).

19 Lupas, A. N. & Bassler, J. Coiled Coils - A Model System for the 21st Century. Trends Biochem. Sci. 42, 130–140 (2017).

20 Woolfson, D. N. The design of coiled-coil structures and assemblies. Adv. Protein Chem. 70, 79–112 (2005).

21 Meinhardt, B., et al. Cardiolipin Stabilizes and Increases Catalytic Efficiency of Carnitine Palmitoyltransferase II and Its Variants S113L, P50H, and Y479F. Int. J. Mol. Sci. 22 (2021).

22 Wai, T. & Langer, T. Mitochondrial Dynamics and Metabolic Regulation. Trends in endocrinology and metabolism: TEM 27, 105–117 (2016).

23 Nan, J. et al. Molecular regulation of mitochondrial dynamics in cardiac disease. Biochimica et biophysica acta. Molecular cell research 1864, 1260–1273 (2017).

24 Angerer, H. et al. Tracing the tail of ubiquinone in mitochondrial complex I. Biochim. Biophys. Acta 1817, 1776–1784 (2012).

25 Zhu, J., Vinothkumar, K. R. & Hirst, J. Structure of mammalian respiratory complex I. Nature 536, 354–358 (2016).

26 Sabbah, H. N. Targeting the Mitochondria in Heart Failure: A Translational Perspective. JACC. Basic to translational science 5, 88–106 (2020).

27 Cogliati, S. et al. Mitochondrial cristae shape determines respiratory chain supercomplexes assembly and respiratory efficiency. Cell 155, 160–171 (2013).

28 Bertero, E. & Maack, C. Metabolic remodelling in heart failure. Nat. Rev. Cardiol. 15, 457–470 (2018).

29 Richter-Dennerlein, R. et al. DNAJC19, a mitochondrial cochaperone associated with cardiomyopathy, forms a complex with prohibitins to regulate cardiolipin remodeling. Cell Metab. 20, 158–171 (2014).

30 Stroud, D. A. et al. Accessory subunits are integral for assembly and function of human mitochondrial complex I. Nature 538, 123–126 (2016).

31 Formosa, L. E., Dibley, M. G., Stroud, D. A. & Ryan, M. T. Building a complex complex: Assembly of mitochondrial respiratory chain complex I. Seminars in cell & developmental biology 76, 154–162 (2018).

32 Karwi, Q. G., Uddin, G. M., Ho, K. L. & Lopaschuk, G. D. Loss of Metabolic Flexibility in the Failing Heart. Front. Cardiovasc. Med. 5, 68 (2018).

33 Diguet, N. et al. Nicotinamide Riboside Preserves Cardiac Function in a Mouse Model of Dilated Cardiomyopathy. Circulation 137, 2256–2273 (2018).

34 Zhou, B. et al. Boosting NAD level suppresses inflammatory activation of PBMCs in heart failure. J. Clin. Invest. 130, 6054–6063 (2020).

35 Martens, C. R. et al. Chronic nicotinamide riboside supplementation is well-tolerated and elevates NAD(+) in healthy middle-aged and older adults. Nat. Commun. 9, 1286 (2018).

36 Pillai, V. B. et al. Exogenous NAD blocks cardiac hypertrophic response via activation of the SIRT3-LKB1-AMP-activated kinase pathway. J. Biol. Chem. 285, 3133–3144 (2010).

37 Koentges, C. et al. SIRT3 deficiency impairs mitochondrial and contractile function in the heart. Basic Res. Cardiol. 110, 36 (2015).

38 Chen, T. et al. Mouse SIRT3 attenuates hypertrophy-related lipid accumulation in the heart through the deacetylation of LCAD. PLoS One 10, e0118909 (2015).

## References

39 Xu, J. Preparation, culture, and immortalization of mouse embryonic fibroblasts. Curr. Protoc. Mol. Biol**. Chapter** 28, Unit 28.21 (2005).

40 Ehler, E., Moore-Morris, T. & Lange, S. Isolation and culture of neonatal mouse cardiomyocytes. J. Vis. Exp. (2013).

41 deAlmeida, A. C., van Oort, R. J. & Wehrens, X. H. Transverse aortic constriction in mice. J. Vis. Exp. (2010).

42 Frezza, C., Cipolat, S. & Scorrano, L. Organelle isolation: functional mitochondria from mouse liver, muscle and cultured fibroblasts. Nat. Protoc. 2, 287–295 (2007).

43 Cox, J. & Mann, M. MaxQuant enables high peptide identification rates, individualized p.p.b.-range mass accuracies and proteome-wide protein quantification. Nat. Biotechnol. 26, 1367–1372 (2008).

44 Webb, B. & Sali, A. Comparative Protein Structure Modeling Using MODELLER. Curr. Protoc. Bioinformatics. 54, 5.6.1–5.6.37 (2016).

45 Maier, J. A. et al. ff14SB: Improving the Accuracy of Protein Side Chain and Backbone Parameters from ff99SB. J. Chem. Theory Comput. 11, 3696–3713 (2015).

46 Wang, J., Wolf, R. M., Caldwell, J. W., Kollman, P. A. & Case, D. A. Development and testing of a general amber force field. J. Comput. Chem. 25, 1157–1174 (2004).

47 Abraham, M. J. et al. GROMACS: High performance molecular simulations through multi-level parallelism from laptops to supercomputers. SoftwareX 1-2, 19–25 (2015).

48 Kabsch, W. & Sander, C. Dictionary of protein secondary structure: pattern recognition of hydrogen-bonded and geometrical features. Biopolymers 22, 2577–2637 (1983).

49 Yan, Y., Tao, H., He, J. & Huang, S. Y. The HDOCK server for integrated protein-protein docking. Nat. Protoc. 15, 1829–1852 (2020).

50 Jo, S., Kim, T., Iyer, V. G. & Im, W. CHARMM-GUI: a web-based graphical user interface for CHARMM. J. Comput. Chem. 29, 1859–1865 (2008).

51 Kumari, R., Kumar, R. & Lynn, A. g_mmpbsa--a GROMACS tool for high-throughput MM-PBSA calculations. J. Chem. Inf. Model. 54, 1951–1962 (2014).

52 Tsugawa, H. et al. MS-DIAL: data-independent MS/MS deconvolution for comprehensive metabolome analysis. Nat. Methods 12, 523–526 (2015).

53 Jewison, T. et al. SMPDB 2.0: big improvements to the Small Molecule Pathway Database. Nucleic Acids Res. 42, D478–484 (2014).

54 Pang, Z. et al. MetaboAnalyst 5.0: narrowing the gap between raw spectra and functional insights. Nucleic Acids Res. 49, W388–w396 (2021).

